# Seasonal polyphenism underlies the origin of a sterile caste in aphids

**DOI:** 10.1101/2022.08.19.501651

**Authors:** Keigo Uematsu, Mayako Kutsukake, Shuji Shigenobu, Man-Miao Yang, Harunobu Shibao, Takema Fukatsu

## Abstract

The origin of a sterile caste among eusocial animals has been a fundamental but still unresolved problem in understanding the evolution of biological complexity. At the origin of a sterile caste, recruitment of pre-existing plasticity may lead to produce physiologically, morphologically and behaviorally distinct caste phenotypes. Here, we provide convincing evidence that preexisting seasonal polyphenism has been recruited to generate a sterile soldier caste in host-alternating social aphids. We demonstrate that sterile soldier nymphs of *Colophina* aphids resemble those of monomorphic defensive nymphs produced in a different host-plant generation. Notably, the two morphs in the basal species show the closest similarity in morphology and gene expression among all morph pairs. Moreover, their evolutionary phenotypic changes along the phylogeny of four *Colophina* species are significantly correlated positively. These results suggest that they may share the common regulatory mechanisms of development, which underpin the heterochronic expression of monomorphic defenders on the different host plant leading to the evolution of a novel soldier phenotype. We further demonstrate that the monomorphic defenders can increase their inclusive fitness by killing predator’s eggs on a seasonally different host plant. Taken together, our findings suggest that preexisting plasticity that can gain indirect fitness benefits facilitates the early evolution of a sterile caste.

## Main text

The origin of sterile castes among eusocial animals has been a fundamental problem in understanding the evolution of biological complexity. (1–4). The sterile caste has morphologically, physiologically, and behaviorally distinct phenotypes from other reproductive individuals despite sharing the same genome and establishing a well-organized society together. Conventionally, the ultimate mechanism underlying the evolution of sterile castes has been argued in the context of kin selection (inclusive fitness) theory (5, 6), but the proximate mechanism underpinning the evolution of sterile castes, especially at the origin of a sterile phenotype, needs additional explanation encompassing the genetic, physiological, and/or developmental changes involved in their caste evolution (7–9).

Developmental plasticity, which generates phenotypic variants in the face of environmental changes, may have facilitated the evolution of novel and complex phenotypes (10–15). Major hypotheses regarding the origin of eusociality suggest that a spatiotemporal regulatory change in a developmental pathway, which stems from preexisting plasticity, has provided an initial caste phenotype. For example, physiological and behavioral differences between non-diapausing and diapausing generations of an ancestral wasp might be recruited to produce worker and queen phenotypes (16, 17), or the decoupling of an ancestral endocrine cycle that regulates the correlated suite of foraging or maternal care behaviors of social bees might produce caste phenotypes (7, 10, 18). These mechanistic hypotheses have been tested by identifying the correlation or similarities between possible ancestral and derived traits using morphological, physiological, and genomic analyses (19–23). However, most recent studies in social insects have been based on the similarity of phenotypes within a single species or a few species from phylogenetically distant groups with different levels of sociality. Such phylogenetically controlled comparisons should include multiple, preferably closely-related species from which possible ancestral states can be inferred in a rigorous manner (24–29).

We investigated the developmental origin of sterile castes in aphids that have a complex, multiple-generation life cycle, during which they seasonally change their host plants and produce morphologically distinct phenotypes through parthenogenesis (30–32). Such seasonal polyphenism, which has evolved multiple times and stems from as far back as the Cretaceous (33, 34), has been considered as an adaptation to various temperatures and different hosts, as well as social interactions such as dispersal, defense, and mimicry (35). Caste differentiation occurs in young (first or second instar) nymphs, called soldiers, that show distinct morphology and behavior for colony defense against predators, and also housekeeping of their nests in some species (36–39). We investigated the soldier caste evolution in aphids of the genus *Colophina* (Eriosomatinae), which shows facultative host alternation between primary host plant (PH), *Zelkova serrata*, and secondary host plants (SH), *Clematis* spp. (Fig. 1A and Fig. 2). On PH, they form a gall, where young first- or second-instar nymphs perform colony defense as monomorphic, non-sterile altruistic defenders (Gall defenders) (40–42). On SH, they form free-living open colonies, where first-instar sterile soldier nymphs (Soldiers) with specialized morphology defend the colony (Fig. 1B-C), whereas first-instar reproductive nymphs (Reproductives) develop and reproduce. This caste differentiation occurs at an early embryonic stage (43). Though appearing at different stages of the life cycle on different host plants, Gall defenders and Soldiers share such characteristic morphological traits as short mouthparts and enlarged forelegs and midlegs (40, 44). While Gall defenders on PH have evolved in multiple lineages of the subfamily Eriosomatinae, Soldiers on SH evolved only in the genus *Colophina* (45–47). The functional and morphological similarities between Soldiers and Gall defenders in *Colophina* has led to the “generation slip hypothesis”, assuming that the traits of non-sterile defenders of the PH generation are heterochronically expressed in the SH generation to predispose the evolution sterile soldier case on SH (48), though this idea has been under controversy (10, 49–51).

**Figure 1.**
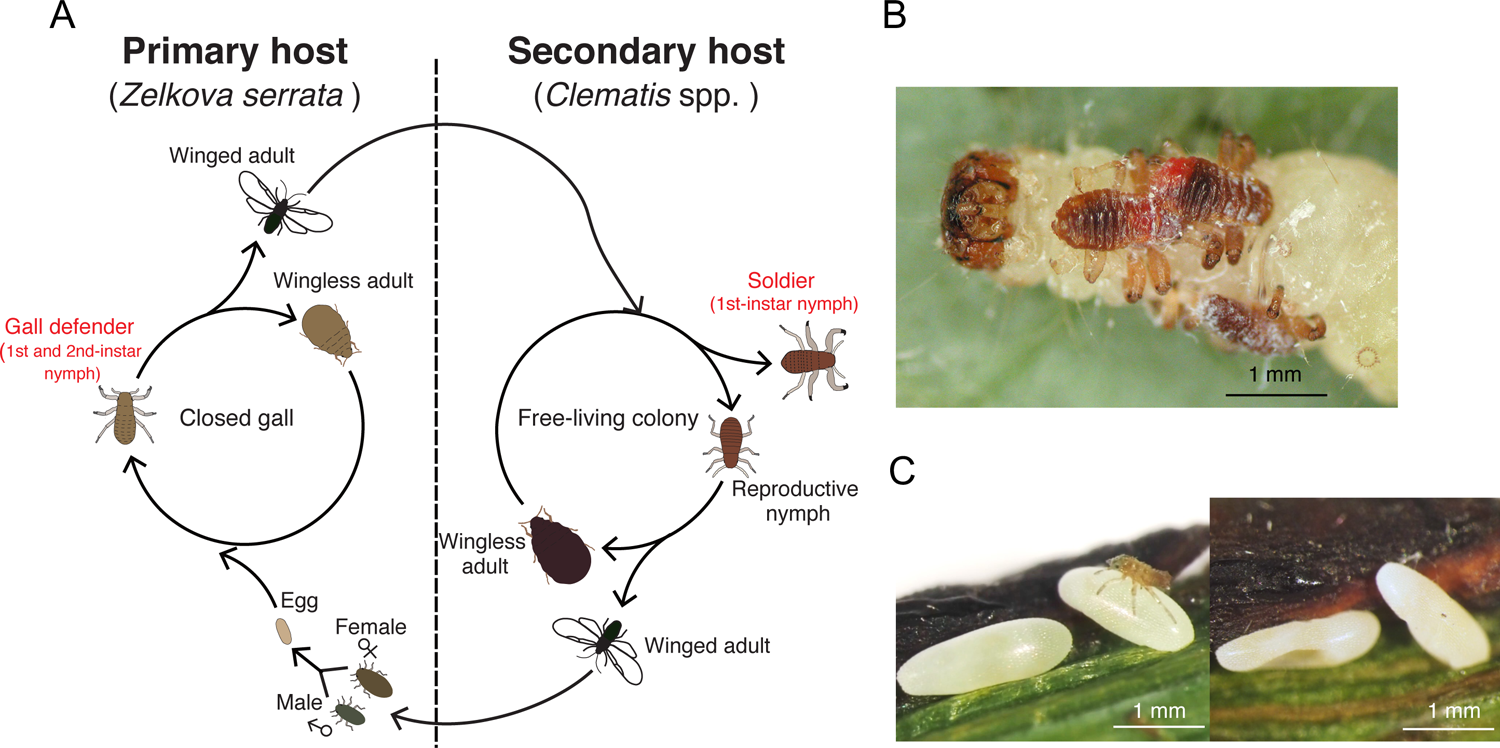
(A) Life cycle of *Colophina* aphids. A foundress hatched from an overwintered egg forms a gall on the leaves of the primary host plant *Zelkova serrata*, wherein young first- or second-instar nymphs (Gall defenders) perform colony defense as monomorphic, non-sterile altruistic defenders. Winged adults migrate to the secondary host plant, *Clematis* spp., and their offspring form woolly colonies, where first-instar sterile soldier nymphs (Soldiers) with specialized morphology defend the colony, and first-instar reproductive nymphs (Reproductives) develop and reproduce. Winged adults flown back to *Z. serrata* and produce males and females, but some nymphs remain on *Clematis* to overwinter and continue clonal reproduction. (B) *C. arma* Soldiers attacking a moth caterpillar. (C) *C. clematicola* Soldiers attacking eggs of a syrphid fly using mouthparts (Left) and the eggs crushed 24 hours after the attack (Right).

**Figure 2.**
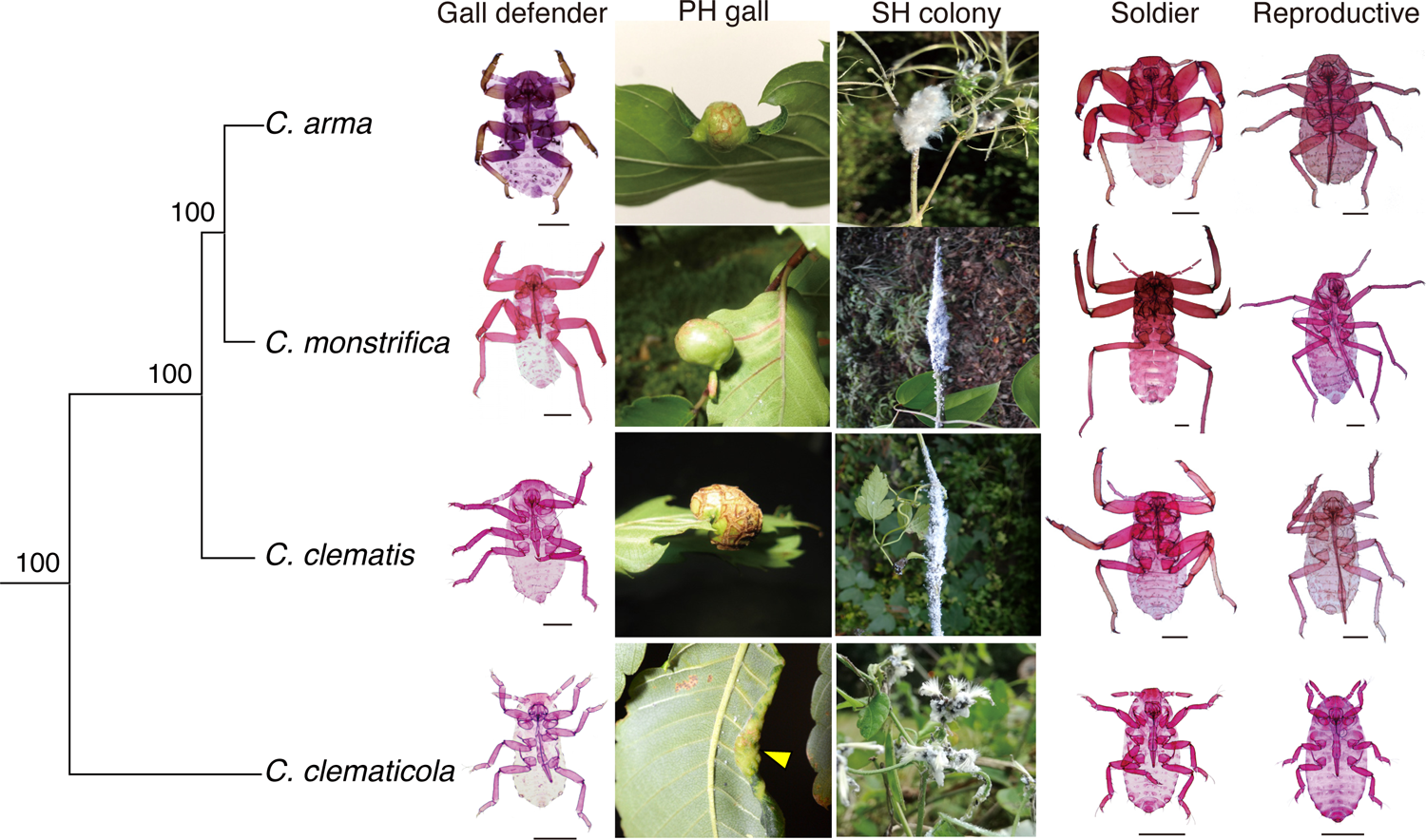
Seasonal and caste polyphenism in first-instar nymphs in *Colophina* aphids and their colonies on the host plants. Shown are Gall defenders and galls on the primary host plant (PH), *Z. serrata*, and Soldiers and Reproductives and free-living colonies on the secondary host plants (SH), *Clematis* spp. of the four *Colophina* species examined in this study. An arrowhead indicates a leaf-rolling gall of *C. clematicola*. Slide-mounted specimens of each species are also shown. The maximum likelihood molecular phylogeny of the four *Colophina* species is generated using 3,755 transcripts that are orthologous among the four *Colophina* species, the apple aphid *Eriosoma lanigerum* and the pea aphid *Acyrthosiphon pisum* (see also Fig. S2). Numbers inside the node indicate bootstrap support values. Soldiers in the derived three species (*C. arma, C. clematis and C. monstrifica*) have enlarged fore- and mid-legs and shortened mouthparts with which to attack predators, whereas Soldiers in the basal species (*C. clematicola*) do not have a well-armored morphology. Scale bars indicate 0.2 mm.

The present study assessed whether the recruitment of preexisting plasticity, seasonal polyphenism, plays an important role in caste evolution among social aphids, by investigating whether a sterile soldier caste (i.e. Soldiers) and its putatively ancestral phenotype (i.e. Gall defenders) share a developmental origin in *Colophina* aphids. The hypothesis that the sterile soldier caste with specialized morphology evolved by recruiting the developmental program of non-sterile Gall defenders predicts a higher level of similarity and correlated evolution between these defensive phenotypes than between other pairs of morphs. Thus, we investigated similarities and evolutionary correlations between Soldiers and Gall defenders through comparisons of closely related *Colophina* species, using morphological, transcriptomic and behavioral data. These investigations provide convincing evidence for the developmental origin of a sterile caste and highlight indirect fitness benefit as a force to fix an original sterile phenotype in the population.

## Results and Discussion

### Correlated evolution of morphological traits

We compared four species of *Colophina* aphids: three species with muscular Soldiers and one species with non-muscular Soldiers. Compared with Reproductives, Soldiers of *C. arma*, *C. clematis* and *C. monstrifica* have enlarged fore- and mid-legs to clutch to predators and a short and robust mouthpart to pierce and kill them (Fig. 1B, Movie S1) (53–56), while Soldiers of *C. clematicola* exhibit no enlargement in legs and sting and destroy eggs of predators by the mouthpart (Fig. 1C, Movie S2. (57, 58)). The non-muscular Soldiers of *C. clematicola* neither molted nor reproduced on SH, *Clematis terniflora* (Table S1), as previously shown by the observation that Soldiers lack next instar skin under their exoskeletons (58). Gall defenders of the three species with muscular Soldiers on SH also had enlarged forelegs, midlegs and short mouthparts to attack predators in the same way as Soldiers (Fig. 2, Movie S3). Molecular phylogenetic analysis based on 3,755 orthologous transcripts showed that the four *Colophina* species formed a monophyletic group in which *C. clematicola* was placed at the basal position (Fig. S2). Given that the muscular Soldiers in the SH generation have only been found in the three *Colophina* species among the Eriosomatinae, the original sterile soldier caste must have a non-muscular form like *C. clematicola*, and the muscular individuals of both PH and SH generations would have evolved after the three species diverged from *C. clematicola*.

To evaluate the morphological similarity between Soldiers and Gall defenders, we measured and analyzed body parts of Soldiers and Reproductives. Analysis of covariance (ANCOVA) revealed that most of the body parts in the three species significantly differed between the two morphs, especially in the enlarged femur and tibia, as well as the shortened tarsus and mouthparts of Soldiers (Fig. 3A and Table S2). We then conducted principal component analysis (PCA) to extract a major variable responsible for caste differentiation (Fig. 3B). The first principal component (PC1; 93.4% contribution) was positively loaded with all traits, indicating the relative sizes of each body part, whereas the second principal component (PC2; 4.7% contribution) was positively loaded with femur and tibia length and negatively loaded with mouthpart and tarsus length (Table S3). This result indicated that the PC2 variable reflects the morphological changes associated with caste differentiation.

**Figure 3.**
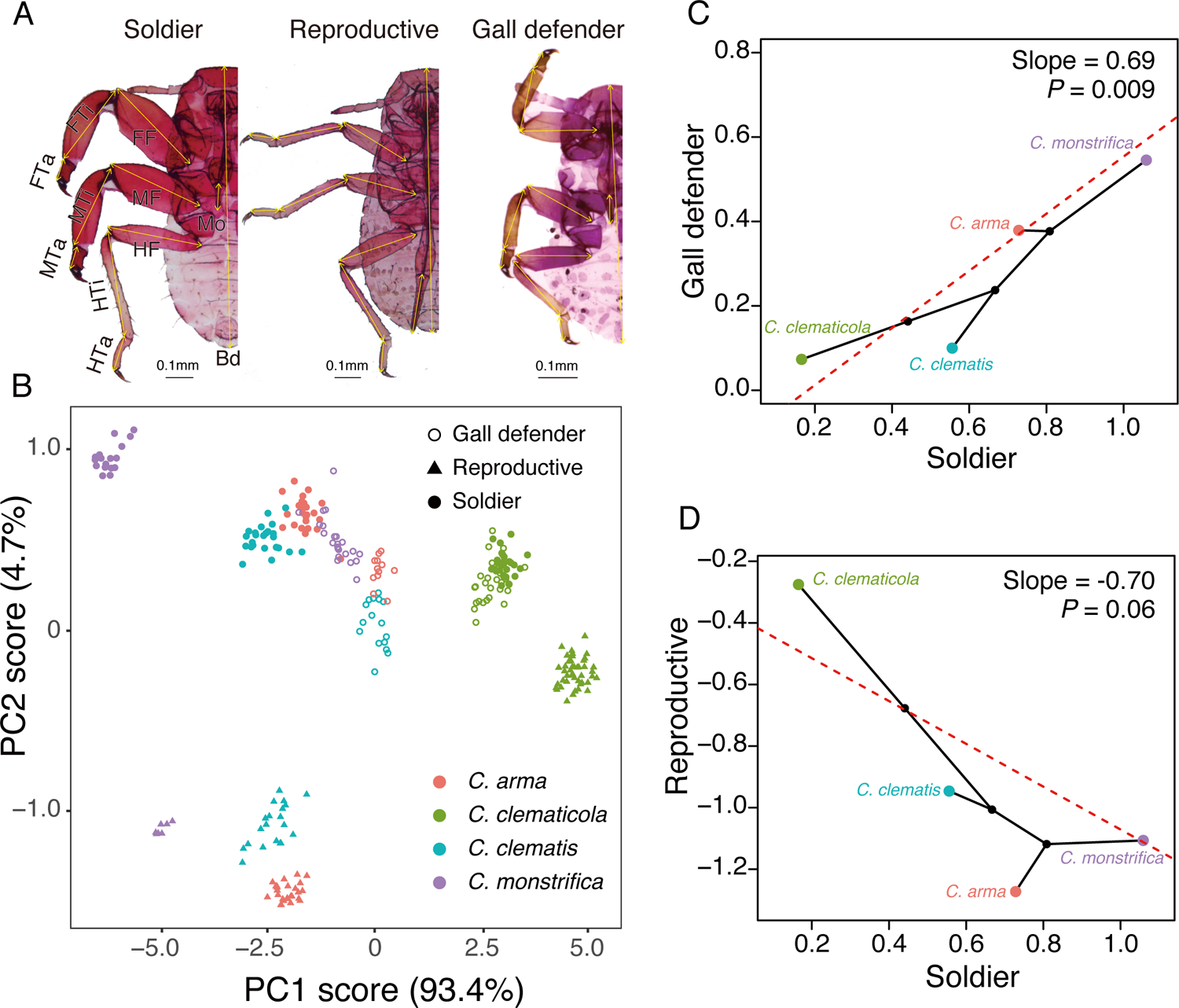
Similarity and correlated evolution in morphological traits between Soldiers on PH and Gall defenders on SH. (A) Three morphs in the first-instar nymphs of *C. arma*: Gall defender (Left); Soldier (Middle); and Reproductive (Right). Morphological characters used in this study are shown. FF, fore femur length; FTi, fore tibia length; FTa, fore tarsus length; MF, mid femur length; MTi, mid tibia length; MTa, mid tarsus length; HF, hind femur length; HTi, hind tibia length; HTa, hind tarsus length; Mo, mouthpart length; Bo, body length. (B) Principal component analysis using 11 morphological characteristics of the first-instar nymphs of the four *Colophina* species. Data point shapes denote morphs: open circles indicate Gall defenders; triangles indicate Reproductives; filled circles indicate Soldiers. See also Table S3 for factor loadings. (C and D) Evolutionary correlation in PC2 scores between (C) Soldiers and Gall defenders and (D) Soldiers and Reproductives. Black filled circles represent internal nodes and their ancestral phenotypic values are estimated by the function *phylomorphospace* in the R package *phytools*. The PC2 score significantly and positively correlated between Gall defenders and Soldiers (*P* = 0.004), while Soldiers and Reproductives showed a negative correlation that is not significant (*P* = 0.062). Colors denote species: *C. arma* = red, *C. clematicola* = green, *C. clematis* = blue, and *C. monstrifica* = purple.

We investigated evolutionary correlations in morphology between Gall defenders and Soldiers by focusing on the PC2 loading scores. We applied phylogenetically controlled regression analysis considering within-species variations according to Ives et al. (59). The PC2 scores significantly and positively correlated between Gall defenders and Soldiers (slope = 0.69, Chi-square = 8.21, df = 1, *P* = 0.004; Fig. 3C). By contrast, the PC2 scores of Soldiers and Reproductives showed a marginally significant negative correlation (slope = −0.70, Chi-square = 3.49, df = 1, *P* = 0.062; Fig. 3D). These results indicate that caste-differentiated traits of Soldiers exhibit higher evolutionary correlation with those of Gall defenders, although they are on different host plants and distinct ecologically and biologically, which suggest the possibility that their defensive traits have been controlled by a shared developmental mechanism.

### Single-species analysis of transcriptomic similarity

To investigate the evolution of gene expression profiles, we performed transcriptomic analysis of the four *Colophina* species. The *de novo* assembled transcriptomes recovered 90.5% – 91.7% of the BUSCO core gene dataset with complete coverage and 5.9% – 6.8% with partial coverage, indicating high completeness of the dataset (Table S4).

Next, the following approaches were applied: single-species analysis using transcriptomes from assembled contigs and then multi-species analysis using orthologs conserved among the four *Colophina* species. First, we identified differentially expressed transcripts (DET) between Soldiers and Reproductives, and DET between Gall defenders and other morphs on PH for each species. The number of significant DET greatly varied depending on species, particularly in Gall defenders. We identified 730, 790, 1,113, and 573 upregulated DET and 685, 957, 1,808, and 1,108 downregulated DET in Soldiers, and 730, 826, 35 and 15 upregulated DET and 202, 255, 84 and 7 downregulated DET in Gall defenders of *C. arma*, *C. clematis*, *C. clematicola*, and *C. monstrifica*, respectively. BLAST and GO analysis showed that Soldier DET shared among at least three species included muscle protein genes such as myosin and troponin, and neurotransmitter-related genes, such as acetylcholine transporter, reflecting muscular morphology and attacking behavior of Soldiers, as well as in the minichromosome maintenance (MCM) complex that controls DNA replication, suggesting presumable suppression of cell proliferation in non-growing Soldiers (Table S5-S6, Dataset S1-S4). Pairwise comparisons between Soldier and Gall defender DET overlapped far more significantly, being 8 – 25–fold higher in upregulated DET and 3 – 10–fold higher in downregulated DET than predicted by chance in all four *Colophina* species, regardless of whether they were upregulated or downregulated (Fig. 4A; *P* < 0.001 in all four species; Fisher exact tests). Single-species hierarchical clustering using all transcripts showed that the expression profiles of these morphs in all four *Colophina* species generally fell into two major classes, which were PH generation and SH generation rather than Soldiers and non-Soldiers (Fig. 4B and Fig. S4). Soldiers clustered with the other morphs on SH in the analysis of the whole transcripts in three species (*C. arma*, *C. clematis* and *C. monstrifica*) that produce soldiers with specialized morphology. By contrast, Soldiers clustered with morphs on PH when only Soldier DET were included (Fig. S4). Notably, in the basal species *C. clematicola*, Soldiers on SH most closely resembled with first-instar nymphs on PH both in the analysis of whole transcripts and Soldier DET, which is remarkable when considering that their emergence seasons and ecological conditions are different (57, 60) (Fig. 4B and Fig. S5). These results indicate that the differentially expressed genes in Soldiers of SH generation resembles the differentially expressed genes in Gall defenders of PH generation, which favor the hypothesis that the Soldier differentiation on SH may share the molecular mechanisms in common with the Gall defender development on PH.

**Figure 4.**
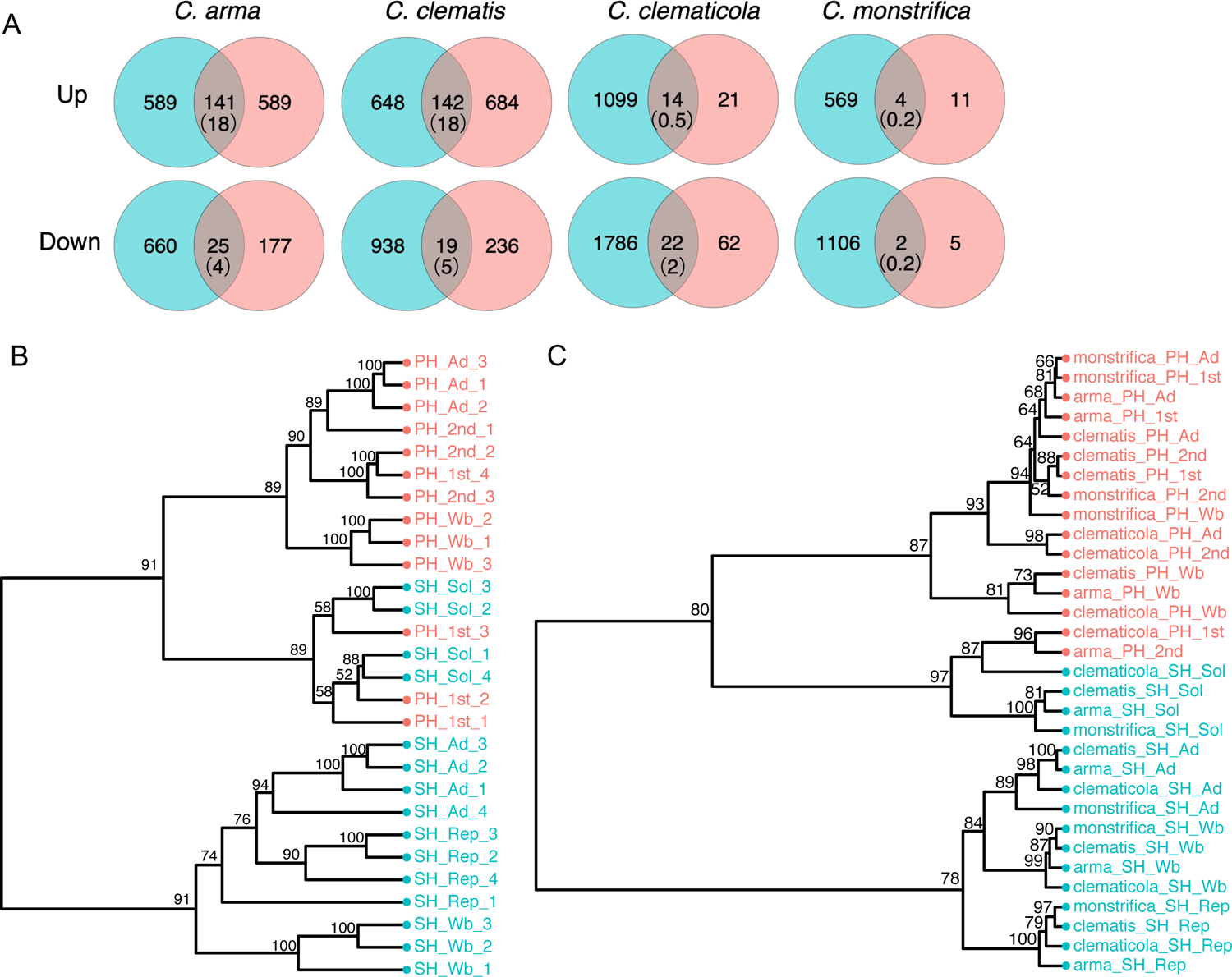
Similarity in transcriptomes between Soldiers and Gall defenders. (A) Venn diagram of the number of differentially upregulated (Upper) and downregulated (Lower) transcripts between Soldiers and Reproductives and between Gall defenders and other morphs in the galls within each species, determined by the R package *edgeR* (FDR-corrected *P* < 0.05). In parentheses are expected numbers of DET if by chance. (B) Transcriptome similarity in *C. clematicola*. Shown is the average relative expression including all transcripts (n = 16,603). Clustering is based on pairwise distance between samples computed as 1 - ρ, where ρ is Spearman’s correlation coefficient. Numbers at nodes represent approximately unbiased multi-scale bootstrap support values. (C) Hierarchical clustering of gene expression levels among four *Colophina* aphids using multi-species Soldier DET across the three *Colophina* species (n = 72). Colors denote host-plant generations: primary hosts (PH), red; secondary hosts (SH), blue. Abbreviations: 1st, first-instar nymph; 2nd, second-instar nymph; Wb, third- and fourth-instar nymph with wing buds; Ad, wingless adults. Hierarchical clustering using all transcripts are indicated in Fig. S5C.

### Multi-species analysis of transcriptomic similarity

We next investigated similarity and correlated evolutionary changes in transcriptomes, using one-to-one orthologous transcripts across the four *Colophina* species. Of 6,287 one-to-one orthologous transcripts, comparative analysis of Soldier DET identified 17 commonly upregulated transcripts and 17 commonly downregulated transcripts across the four species, whose occurrences were 1119– and 686–fold higher than those predicted by chance (0.015 and 0.025; Fig. S6A and B; *P* < 0.001 for both cases; Fisher exact tests). In addition, a similar level of overlaps in Soldier DET between *C. clematicola* and the other three species substantiated the notion that they share a molecular basis for caste differentiation, indicating the single origin of the soldier caste despite differences in their morphology in relation to colony defense.

We selected 6,017 transcripts that contained orthologous sequences longer than 200 bp across the four species and re-mapping raw reads to them. Multi-species hierarchical clustering showed that the transcriptomes clustered primarily by species and that Soldiers were not grouped with other phenotypes when all orthologs were included (Fig. S6C). By contrast, Soldiers grouped with the morphs on PH, but not with those on SH when 72 orthologous Soldier DET across the three derived species were included (Fig. 4C). These results indicate that the resemblance of gene expression involved in soldier differentiation on SH has been conserved since the common ancestor of *Colophina*, suggesting that their caste development has been controlled by a molecular mechanism that concurrently controls the gene expression of the seasonally different PH generation.

### Correlated evolution of transcriptomes

We estimated levels of correlated evolution by calculating correlation coefficients of PIC (61) of the 6,017 orthologous transcripts and compared evolutionary changes in expression levels within a phylogenetic context, as used in recent studies to show homology between body regions within an individual (29). Soldiers on SH had higher Pearson average correlation values in the PIC including all orthologous transcripts with Gall defenders on PH (*r* = 0.48 with first-instar Gall defenders; 0.49 with second instar Gall defenders) than with Reproductives on SH (*r* = 0.37) (Fig. S7, Table S7), showing a significant difference (permutation tests, first-instar Gall defenders vs. Reproductives: *P* = 0.006; second-instar Gall defenders vs. Reproductives: *P* = 0.009, Fig. S8). In particular, Soldiers had highest correlation values with Gall defenders for multi-species Soldier DET (*r* = 0.52 with first-instar Gall defenders; 0.55 with second instar Gall defenders, Fig. 5A and 5C, Table S8) among the pairs, whereas Soldiers and Reproductives had lower correlation coefficients (*r* = 0.18, Fig. 5B), showing a significant difference (permutation tests, first-instar Gall defenders vs. Reproductives: *P* = 0.003; second-instar Gall defenders vs. Reproductives: *P* = 0.003, Fig. S8).

**Figure 5.**
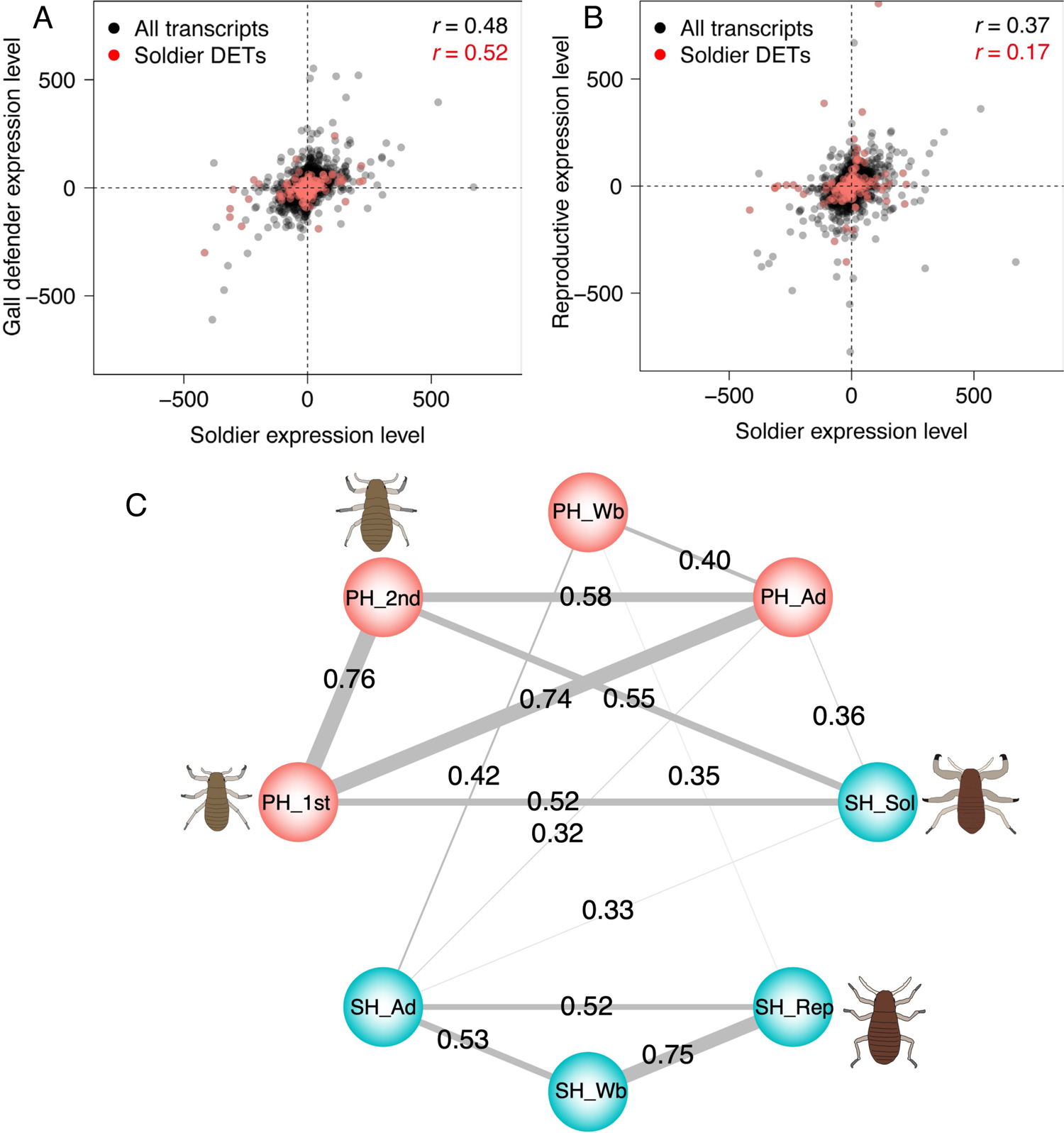
Correlated evolution in transcriptomes between Soldiers and Gall defenders. (A) and (B) Scatter plots of phylogenetic independent contrasts (PIC) for gene expression levels, including all pairwise three branches (n = 18,051 shown in black for all transcripts and n = 216 shown in red for Soldier DET). (C) Correlation networks using PIC in the Soldier DET. A number on the line represents Pearson’s Pairwise correlation coefficient (*r*) between a pair of morphs. Only *r* > 0.3 is shown as a line. Thicker line indicates a higher correlation. In (C), Color coding and abbreviations are used as in Fig. 4.

Overall, despite different emergence seasons and ecological conditions, the whole-body transcriptomes of sterile Soldiers and non-sterile Gall defenders on different host plants were similar and evolutionarily correlated, especially with respect to transcripts associated with caste differentiation. This high degree of evolutionary correlation shows that the gene expression levels in the two defensive nymphs have changed in a correlated manner since the common ancestor of *Colophina*, suggesting that they have shared a regulatory mechanism after the evolution of Soldiers. Therefore, taken together with the morphological findings described above, the phenotypes of Soldiers and Gall defenders may have shared a developmental origin, diverged after the evolution of a soldier caste and been adapted to different ecological conditions within their life cycle. Their evolutionary relationship is reminiscent of serially homologous characters within the same organism, such as hands and feet in primates (27, 28).

### Effective colony defense on a seasonally different host plant in the basal species

We investigated the phenotype and its fitness benefits at the origin of eusociality by analyzing the basal species *C. clematicola* more closely as a proxy of the ancestral eusocial species. Soldiers and Gall defenders of *C. clematicola* are very similar morphologically (Fig. S9), which is also shown by PCA (Fig. 3B), and shared many characteristics, such as the presence of cornicles (61) or longer leg parts (Fig. S10). In addition, clustering analysis of transcriptomes showed that Soldiers and Gall defenders were the most closely related within the species (Fig. 4B and Fig. S6).

The high levels of morphological and transcriptomic similarity between Soldiers and Gall defenders of *C. clematicola* prompted us to examine their similarities in behavior related to colony defense. In the derived three species, *C. arma*, *C. clematis*, and *C. monstrifica*, Soldiers and Gall defenders are known to show similar attacking behavior by clutching predators using enlarged forelegs and midlegs and piercing them using shortened mouthparts (Movie S1 and S3). In *C. clematicola*, soldiers show egg-destroying defensive behavior using mouthparts, instead of attacking insects (Fig. 1C and Movie S2). Although egg-destroying behavior of Gall defenders of *C. clematicola* has not been identified (61), their closely similar morphology and transcriptomes predicted that Gall defenders would also exhibit the same defense behaviors as Soldiers. Indeed, we found that Gall defenders showed egg-destroying behavior by inserting stylets into eggs for more than 15 minutes (Fig. 6). We tested the effects of defensive behaviors of them on different host plants using eggs from syrphid flies that are the common predators of *Colophina* species on SH (62–64), and found that their eggs are actually attacked by *C. clematicola* Soldiers (57). 17 (74%) of 23 syrphid eggs attacked by Gall defenders and 12 (100%) of 12 attacked by Soldiers were significantly deformed and eventually unhatched in contrast to no or a few unhatched eggs in controls (*P* < 0.01, Fisher exact tests, Fig. 6). These results indicate that Gall defenders are able to effectively defend free-living colonies on SH. Note that syrphid flies are not the main predators of the PH generation, as 38 (50%) of 76 of *Colophina clematis* galls collected from three distant field sites were invaded by the aphid fly *Leucopis* spp. (Table S9). These flies laid eggs around the galls and the hatched larvae preyed on the aphids inside (Fig. S11). In contrast, only 1 (1.3%) of 76 galls was invaded by a syrphid fly larva (*P* < 0.001, Fisher exact test). This indicates that Gall defenders of *C. clematicola* can increase their indirect fitness by destroying the eggs of predators, even if their phenotypes are accidentally expressed on a seasonally and ecologically different host plant that is unsuitable for their growth and reproduction.

**Figure 6.**
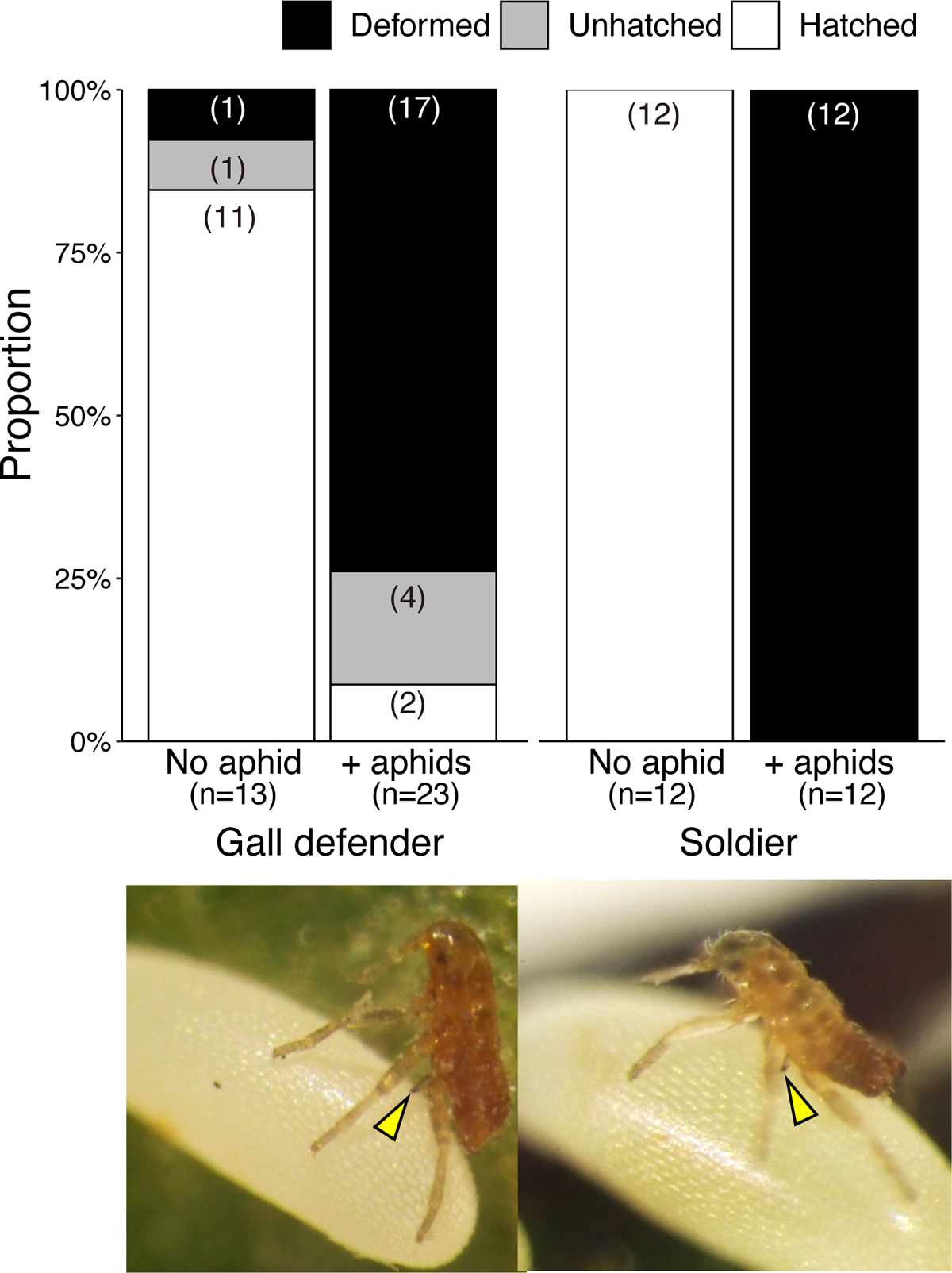
Close phenotypic resemblance between Soldiers and Gall defenders in the basal species *C. clematicola*. The fate of predator eggs attacked by Soldiers and Gall defenders are shown. Pictures below indicate corresponding nymphs of *C. clematicola* attacking predator eggs. Arrowheads indicate stylets injected into the predator egg.

### Heterochronic expression of a seasonally different host-plant generation explains a developmental origin of a sterile caste

Our results provide lines of evidence that the non-sterile Gall defenders and the sterile Soldiers, produced in different seasons and ecological conditions, shared a regulatory mechanism that has been conserved since the origin of a sterile soldier caste. They share morphologies and gene expression that are differentiated during caste development (Fig. 3B, 4A and 4C). In addition, phylogeny-controlled comparisons among four *Colophina* species found that the evolutionary correlation between the two defensive morphs is closer than that between Soldiers and Reproductives or other morphs of SH generations (Fig. 3C-D, 5A-C). Furthermore, we found that the similarity of the two defensive morphs in the basal lineage was the highest among all pairwise morph comparisons (Fig. 3B and 4B), despite different ecological conditions between the two host-plant generations. Thus, these results provide quantitative evidence for the hypothesis that a sterile soldier caste on SH evolved by recruiting the developmental program of the non-sterile Gall defenders on PH. Heterochronic expression of phenotype on a different host-plant generation has been occasionally found in some aphid species, including Eriosomatini, and its importance in the evolution of the aphid complex life cycle has been suggested (30, 48, 49, 65). Although its underlying molecular mechanism remains to be solved, a similar heterochronic expression of the Gall defender phenotype on SH likely contributes to the recruitment of seasonal polyphenism and evolution of a novel soldier caste in aphids, and the subsequent divergent selection might shape the specialized morphology of Soldiers for attacking predators found in the derived species.

Previous findings of other social insects have suggested that plasticity in an ancestral life cycle was recruited to the novel phenotype of a sterile caste (16, 17, 19, 20, 66), but evidence was based on comparisons of morphs within a single species or distant clades, and thus its detailed evolutionary process has been largely unknown. In *Colophina* aphids, by contrast, the putative ancestral phenotype, Gall defender, persists within their life cycle, thus correlations in the two evolutionary trajectories between the ancestral and derived phenotypes can be investigated quantitatively using multiple species in the same genus (Fig. 3C, 4C and D). Thus, our results provide unprecedentedly detailed evidence for a shared molecular mechanism that pleiotropically regulates the development of sterile caste and its putatively ancestral phenotype, suggesting plasticity-based caste evolution in social insects.

### Indirect fitness effects at the early stage of heterochrony-based caste evolution

Our behavioral experiment in the basal species *C. clematicola* provides insight into the origin of a sterile soldier caste in aphids. Aphids are morphologically and physiologically specialized to their two host plants in their life cycle; thus, the original altruistic defenders on SH with a phenotype closely resembling that on PH should have had difficulty in growing and reproducing by feeding on sap produced by different host plants. However, such a decrease in direct fitness can be compensated by the defensive phenotype that increases indirect fitness through colony defense such as attacking predators or destroying their eggs, as shown in *C. clematicola* (Fig. 5B). Thus, our results suggest the possibility that kin selection may have facilitated the early evolution of a novel caste phenotype expressed in a seasonally and ecologically different environment through heterochronic change.

### Concluding remarks

Overall, our study provides strong evidence that recruiting a preexisting seasonal polyphenism played an important role in the evolution of the novel phenotype of a sterile soldier caste in aphids. Future experimental investigations of *Colophina* and its related non-soldier-producing species under controlled laboratory conditions should reveal which types of genetic or environmental changes caused the heterochronic expression of Gall defenders on SH and thus reveal the actual processes involved in the plasticity-based evolution of a novel phenotype.

## Materials and methods

### Sample collection

Galls on *Z. serrata* trees and free-living colonies on *Clematis* plants of *Colophina* aphids were collected under natural conditions as follows: *C. arma* in Akiota, Hiroshima; *C. clematicola* in Tsukuba, Ibaraki; *C. clematis* in Shomaru, Saitama, Japan and *C. monstrifica* in Huisun Forest, Nantou, Taiwan (Table S10). Galls of *C. clematicola* were artificially induced (see below) on *Z. serrata* because they have been never found in the field. After a series of experiments, the aphids were collected and brought to laboratories for further analyses.

### Artificial induction of gall generations of *C. clematicola*

Winged adults (sexuparae) of *C. clematicola* were collected from *Clematis terniflora* (AIST, Tsukuba, Japan) and maintained outdoors in clear plastic containers with bundles of *Zelkova serrata* bark to encourage the production of male and female sexual nymphs as described (61). The sexual nymphs can reach adulthood without feeding, copulate and produce eggs. From April, the containers were monitored daily and gall foundress nymphs that hatched were attached to shooting branches of *Z. serrata* trees using a fine brush. Induced galls by the gall foundress nymphs were covered with a bag to protect against predators until collection for laboratory experiments.

### Morphometry

Aphids were preserved in 70% ethanol, boiled in 10% KOH, stained with acidic fuchsin, dehydrated through an ethanol-xylene series, mounted on microscope slides in balsam or Mount-Quick (Daido Sangyo, Saitama, Japan). The mounted individuals were photographed using a Nikon 1 digital camera (Nikon, Tokyo, Japan) attached to a light microscope. The morphological characteristics of the length of the fore, mid and hind femur, tibia and tarsus, as well as the ultimate rostral segments as mouthparts and the body were measured using ImageJ. Landmarks used for measurement are shown in Fig. S3. The data were log-transformed to satisfy normality and constant variance and then compared using analysis of covariance (ANCOVA) using body length as a covariable. All data were statistically analyzed using R version 3.3.3 software.

### Library preparation and sequencing

Total RNA was extracted from whole bodies of the four aphid species preserved in acetone. We collected first- (n = 10 – 20 pooled individuals) and second- (n = 5 – 10) instar nymphs, third- or fourth-instar nymphs with wing buds (n = 1 or 2) and wingless adults (n = 1 or 2) from the PH generation, and first-instar soldier and reproductive nymphs, third- or fourth-instar nymphs with wing buds and wingless adults from the SH generation. For each species, three or four biological replicates are prepared for each morph, except that seven replicates prepared for the PH generation of *C. clematis*. We extracted total RNA using RNeasy (Qiagen, Hilden, Germany) column purification kits or Maxwell 16 LEV simplyRNA Purification Kits (Promega, Madison, WI, USA) and prepared cDNA libraries using TruSeq RNA Library Prep Kit (Illumina, San Diego, CA, USA). The quality of the libraries was inspected by an Agilent Bioanalyzer 2100 (Agilent Technologies). Samples were separately tagged and sequenced on a HiSeq or NovaSeq (Illumina). We filtered read pairs that contained Illumina adaptor sequences with Trimmomatic v. 0.36 (67) (parameters LEADING:30 TRAILING:30 MINLEN:70), and ribosomal RNA and mitochondria DNA.

### Transcriptome assembly, estimation of transcript abundance and differential expression analysis

For each species, *de novo* transcriptome assembly was conducted using Trinity (68) with default parameters. All reads were mapped to Trinity contigs using RSEM (69). For each contig cluster, the isoform with the highest expression level was selected, through a minimum expression filter of two transcripts per million mapped reads (TPM) in at least half of the morphs from either host-plant generation to reduce non-expressed contigs. (Fig. S1). Transcripts were functionally annotated using Trinotate v3.0.1 (https://trinotate.github.io). For each species, the read count estimates were input into the R package *edgeR* (70) to identify differentially expressed transcripts in pairwise morph comparisons using a false discovery rate (FDR) (71) adjusted p-value of 0.05. Each count estimate was normalized using the TMM method in *edgeR*. The normalized count estimates were used to derive a Euclidean distance matrix for clustering analysis using *hclust* function in R. Approximately unbiased p-values were calculated using the R package *pvclust* (72), with bootstrap resampling (1,000 replicates). Differentially expressed transcripts (DET) with at least a 2-fold change in expression were further analyzed. We determined whether the DET expressed within each species pair and tissues exceeded the number of overlapping transcripts using the R package *SuperExactTest* v. 0.99.4 (73) in R, which calculates the probability of multi-set intersections. We used an FDR adjusted p-value of 0.05 in analyses of multiple intersections.

### Cross-species transcriptome comparison

Putative orthologues among the four *Colophina* species were identified using the Basic Local Alignment Search Tool (BLAST) reciprocal best hits in which one-to-one orthologs were selected based on a combination of the lowest e-value and bit-score generated by BLASTn, with an e-value cutoff of 1e^−5^ and a minimum identity of 30%. This approach identified 6,287 orthologous transcripts for the four species. Sequences of the orthologous transcripts were aligned using MAFFT (74) and their terminal gaps were removed using trimAl (75). We selected 6,017 transcripts longer than 200 bp as a reference from the aligned orthologous sequences, then RNA-Seq reads were mapped again to the transcripts to calculate read counts for each species by RSEM. The count estimate was normalized by sqrt (TPM), as used in previous cross-species comparisons. The clustering analysis and bootstrap resampling for multi-species analyses were performed as described in the single-species analysis using the normalized count estimates of all orthologous transcripts or 72 multi-species Soldier DET across the three *Colophina* species (*C. arma, C. clematis,* and *C. monstrifica*) that produce soldiers with specialized morphology.

### Phylogeny

The 6,287 orthologous transcripts among *Colophina* were also examined for orthology with the apple woolly aphid *Eriosoma lanigerum* (76) and pea aphid *Acyrthosiphon pisum* (77) available from AphidBase (https://bipaa.genouest.org/is/aphidbase/) by using reciprocal best BLAST hit. The remaining 3,755 one-to-one orthologous transcripts were aligned using MAFFT (74) and a maximum likelihood phylogeny of the six species was constructed using RaxML (78).

### Statistical analysis for testing correlated evolution

We applied principal component analysis using the R package *prcomp* to summarize the overall morphological changes involved in caste differentiation. The morphological data were log-transformed to satisfy normality and constant variance. Principal components were initially estimated for each species using mean values to avoid the effects of different sample sizes per species, then calculated the first and the second principal component (PC1 and PC2) scores for each individual. We tested for correlations in PC2 values for morphological characteristics between Soldiers and Gall defenders using phylogenetic generalized least-squares (PGLS) analyses using *pgls.Ives* function in the R package *phytools* (79) that account for within-species variation in both variables (59). We used molecular phylogeny estimated from the Because model convergence is often sensitive to starting conditions in *pgls.Ives*, we repeated the function 1,000 times and selected the top 10 models that showed convergence with the highest likelihood, and calculated the mean intercept, slope, and likelihood to set an average model (80). To test for correlations in transcriptomes, we averaged sqrt (TPM) values across replicates to generate a representative transcriptome for each morph in each species and then calculated phylogenetic independent contrasts (PIC) between all morph pairs for the expression levels of each of the 6,017 orthologous transcripts. Pearson correlation coefficients of PIC were calculated using all orthologous transcripts or 72 multi-species Soldier DET across the three *Colophina* species (*C. arma, C. clematis,* and *C. monstrifica*) that produce soldiers with specialized morphology. To test for differences in correlation coefficients between two morph pairs, *r*_1_ – *r*_2_, we randomly drew, without replacement, 10,000 samples of permutated PIC vectors of the same lengths from the two groups and recalculated the difference *r’*_1_ – *r’*_2_ for each. Two-tailed p-values were calculated as the proportion of times the permutated difference exceeded the observed value. The correlation matrix among morphs was visualized as a network using the R package *qgraph* (81).

### Attacks against predator eggs by first-instar nymphs of *Colophina clematicola*

The eggs of syrphid flies collected at Meguro, Tokyo, Japan were placed in plastic Petri dishes (diameter, 35 mm; height 10 mm), then 10 first-instar nymphs of *C. clematicola* were introduced into the dishes. Aggressive behavior was observed for 15 minutes under a dissecting microscope. When they clutched the eggs and their antennae stopped moving for more than 30 seconds, we regarded them as attacking and started measuring how long the nymphs stayed on the eggs as a proxy of injection time. We then individually placed attacked and unharmed eggs in 1.5-mL microcentrifuge tubes and examined their fates daily for seven days.

### Availability of supporting data

Raw RNA-Seq data in FASTQ format were deposited in the DDBJ database under the BioProject accession number PRJDB12646.

## Acknowledgments

We thank Shigeyuki Aoki, Utako Kurosu and Shin-ichi Akimoto for providing slided specimens of *Colophina* aphids and information for their collecting sites. This study was supported by the Sumitomo Foundation and Japan Society for the Promotion of Science to K. U., and NIBB Collaborative Research Program (14-701, 15-803, 16-402) to T. F. Computational resources were provided by the Data Integration and Analysis Facility, National Institute for Basic Biology. The collection of aphids at Huisun Experimental Forest Station was permitted by the Experimental Forest Management Office, National Chung Hsing University (nos. 1020000228 and 1030000107).

## Supporting Information

### Materials and Methods

#### Laboratory rearing the open-colony generation of *Colophina clematicola*

Developing shoots of *Clematis terniflora* with a terminal bud were cut and placed in 6-well plastic tissue culture plates (Iwaki, diameter, 34.6 mm; height, 17.5 mm). Cotton wool soaked with distilled water was then wrapped around the cut ends. Four first-instar nymphs (Soldiers or Reproductives) collected from natural colonies on *Clematis terniflora* at AIST, Tsukuba, Japan were introduced into each well, then the plates were sealed with Parafilm and placed in an incubator at 20°C (12L:12D). The Parafilm and cotton wool were replaced every two days, when the number of molted nymphs and survival rates were recorded.

#### Functional annotation and analysis

Transcripts were functionally annotated using Trinotate v3.0.1 (https://trinotate.github.io). Similarities to function-known proteins were detected using a BLASTx search (e-value ≤ 1e^−5^) of the Swiss-Prot protein database. Likely coding regions within transcripts were predicted using TransDecoder (http://transdecoder.github.io), and resulting protein products were searched for sequence similarities using a BLASTp search (e-value ≤ 1e^−5^) and for conserved protein domains using Hmmer (http://hmmer.org/) and PFam (1). Signal peptides were predicted using SignalP (2), ribosomal RNA was predicted using RNAmmer, and transmembrane regions were predicted using TmHMM (3). We applied Gene Ontology (GO) enrichment analysis to each species to determine whether particular classes of soldier-biased transcripts were enriched for specific functional characteristics. Orthologs of each gene were determined using UniProtKB/Swiss-Prot. The GO term enrichment analysis was conducted using R package *GOseq* (4) with a false discovery rate (FDR)-corrected significance threshold of 0.05.

---

**Table S1.** Molting rate of the first-instar soldier and reproductive nymphs of *C. clematicola*.

|  | Prop. of colonies<br>without molting nymphs<br>after 10 days | Prop. of surviving<br>aphids<br>after 7 days | Prop.of surviving<br>aphids<br>after 10 days | Prop. of molting<br>aphids<br>within 10 days |
| --- | --- | --- | --- | --- |
| Soldier | 100<sup>a</sup> % (17/17) | 13<sup>b</sup> % (9/68) | 0<sup>c</sup> % (0/68) | 0<sup>d</sup> % (0/68) |
| Reproductive | 18<sup>a</sup> % (3/17) | 44<sup>b</sup> % (30/68) | 44<sup>c</sup> % (30/68) | 47<sup>d</sup> % (32/68) |

<sup>abcd</sup>Values within a column with different superscripts are significantly different (Fisher's exact tests, $P < 0.001$ ).

**Table S2.** Morphometric analyses of 10 body measurements in first instar nymphs. The data were log transformed and then compared using analysis of covariance (ANCOVA) using body length as a covariable. Mean values ± SD (μm) are shown.

| Species | *C. arma* |  |  |  |  | *C. clematis* |  |  |  |  |  |  |
| --- | --- | --- | --- | --- | --- | --- | --- | --- | --- | --- | --- | --- |
| Body characteristics | Soldier (N=25) |  | Reproductive (N=26) |  | *F* | *P*-value | Soldier (N=24) |  | Reproductive (N=20) |  | *F* | *P*-value |
|  | Mean | SD | Mean | SD |  |  | Mean | SD | Mean | SD |  |  |
| Fore femur | 359.7 | 19.8 | 282.6 | 11.0 | 604.3 | 7.55E-29 | 387.3 | 23.6 | 300.5 | 16.0 | 264.4 | 1.76E-19 |
| Fore tibia | 296.3 | 15.4 | 265.1 | 13.0 | 201.1 | 8.69E-19 | 351.5 | 21.6 | 296.4 | 18.9 | 115.3 | 1.74E-13 |
| Fore tarsus | 84.8 | 4.0 | 97.4 | 4.1 | 79.9 | 8.77E-12 | 100.3 | 6.0 | 103.8 | 6.4 | 0.1 | 0.74 |
| Mid femur | 333.4 | 18.6 | 280.1 | 9.6 | 367.1 | 3.97E-24 | 381.6 | 20.7 | 296.7 | 16.5 | 316.6 | 6.86E-21 |
| Mid tibia | 301.0 | 14.2 | 287.6 | 13.6 | 65.1 | 1.74E-10 | 359.0 | 20.7 | 316.6 | 20.9 | 68.3 | 2.86E-10 |
| Mid tarsus | 90.2 | 4.3 | 107.7 | 4.2 | 162.4 | 5.10E-17 | 107.2 | 6.4 | 113.0 | 6.8 | 2.2 | 0.15 |
| Hind femur | 323.9 | 19.0 | 305.7 | 11.7 | 88.7 | 1.74E-12 | 361.6 | 18.7 | 316.8 | 17.5 | 99.7 | 1.53E-12 |
| Hind tibia | 359.3 | 21.8 | 333.4 | 16.4 | 69.9 | 6.26E-11 | 425.2 | 24.3 | 364.4 | 22.7 | 106.3 | 5.98E-13 |
| Hind tarsus | 118.9 | 6.5 | 118.9 | 4.0 | 3.4 | 0.07 | 142.5 | 8.5 | 128.5 | 8.2 | 54.1 | 5.23E-09 |
| Ultimate rostral segments | 107.2 | 3.1 | 218.6 | 5.1 | 8389.9 | 1.24E-54 | 119.3 | 4.3 | 195.1 | 11.1 | 1217.8 | 4.10E-32 |

| Species | *C. monstifica* |  |  |  |  | *C. clematicola* |  |  |  |  |  |  |
| --- | --- | --- | --- | --- | --- | --- | --- | --- | --- | --- | --- | --- |
| Body characteristics | Soldier (N=17) |  | Reproductive (N=7) |  | *F* | *P*-value | Soldier (N=25) |  | Reproductive (N=44) |  | *F* | *P*-value |
|  | Mean | SD | Mean | SD |  |  | Mean | SD | Mean | SD |  |  |
| Fore femur | 721.3 | 30.3 | 414.5 | 10.2 | 265.5 | 2.16E-13 | 122.5 | 4.4 | 91.9 | 4.5 | 855.5 | 1.67E-39 |
| Fore tibia | 685.7 | 26.2 | 442.8 | 9.2 | 190.1 | 5.39E-12 | 104.2 | 3.4 | 76.9 | 4.3 | 620.3 | 2.83E-35 |
| Fore tarsus | 144.3 | 8.0 | 165.4 | 6.9 | 50.3 | 5.34E-07 | 58.6 | 2.2 | 41.7 | 1.7 | 1173.6 | 9.31E-44 |
| Mid femur | 714.4 | 28.0 | 409.7 | 10.9 | 272.1 | 1.70E-13 | 126.5 | 4.7 | 90.8 | 4.0 | 1244.2 | 1.49E-44 |
| Mid tibia | 708.0 | 29.7 | 477.0 | 12.8 | 159.0 | 2.90E-11 | 112.7 | 4.6 | 79.5 | 4.0 | 881.5 | 6.66E-40 |
| Mid tarsus | 157.6 | 7.7 | 192.9 | 8.1 | 135.8 | 1.25E-10 | 63.8 | 2.6 | 45.0 | 1.9 | 1183.3 | 7.19E-44 |
| Hind femur | 687.0 | 25.8 | 446.2 | 16.6 | 94.3 | 3.23E-09 | 139.1 | 5.3 | 105.2 | 4.5 | 894.7 | 4.23E-40 |
| Hind tibia | 873.6 | 32.7 | 539.2 | 23.4 | 122.1 | 3.28E-10 | 136.7 | 5.8 | 101.1 | 4.9 | 855.5 | 1.67E-39 |
| Hind tarsus | 269.8 | 9.9 | 210.9 | 3.9 | 26.5 | 4.28E-05 | 73.1 | 2.0 | 53.5 | 2.7 | 904.0 | 3.07E-40 |
| Ultimate rostral segments | 136.5 | 5.2 | 221.8 | 4.5 | 364.5 | 9.49E-15 | 74.5 | 1.9 | 82.9 | 2.8 | 159.2 | 2.98E-19 |

**Table S3.** Factor loadings for principal component analysis.

| Trait | PC1<br>(93.4%) | PC2<br>(4.7%) | PC3<br>(1.3%) | PC4<br>(0.4%) |
| --- | --- | --- | --- | --- |
| Fore femur length | -0.3071 | 0.1976 | -0.1923 | -0.3088 |
| Fore tibia length | -0.3097 | 0.1266 | -0.1248 | -0.2084 |
| Fore tarsus length | -0.3044 | -0.0917 | 0.5314 | 0.0201 |
| Mid femur length | -0.3082 | 0.1876 | -0.1149 | -0.2500 |
| Mid tibia length | -0.3109 | 0.0869 | -0.0478 | -0.1758 |
| Mid tarsus length | -0.3021 | -0.1346 | 0.5859 | 0.1553 |
| Hind femur length | -0.3105 | 0.1090 | -0.0964 | -0.0933 |
| Hind tibia length | -0.3104 | 0.1176 | -0.1106 | -0.0674 |
| Hind tarsus length | -0.3070 | 0.1548 | 0.1882 | 0.3569 |
| Mouthpart length | -0.2359 | -0.9064 | -0.1884 | -0.2042 |
| Body size | -0.3025 | -0.0666 | -0.4624 | 0.7490 |

**Table S4.** RNA sequencing and de novo transcriptome assembly of four *Colophina* species.

| Species | No. of<br>library<br>samples | No. of total<br>raw read<br>pairs | No. of total<br>assembled<br>bases | Prop. of<br>GC<br>content | Prop. of<br>complete<br>BUSCOs<br>(arthopoda_<br>odb10) | Prop.of<br>partial<br>BUSCOs<br>(arthopoda_<br>odb10) | Average<br>number of<br>aligned reads<br>per sample<br>(range) | DRR accession<br>nos. |
| --- | --- | --- | --- | --- | --- | --- | --- | --- |
| *C. arma* | 24 | 287 million | 2.38 million | 31.0% | 91.1%<br>(971/1066) | 6.3%<br>(67/1066) | 8.7 million<br>(5.2-13.3) | DRR346959-346982 |
| *C. clematicola* | 28 | 224 million | 3.25 million | 30.5% | 90.5%<br>(965/1066) | 6.8%<br>(73/1066) | 6.3 million<br>(1.3-10.2) | DRR347007-347028,<br>DRR347053-347058 |
| *C. clematis* | 40 | 375 million | 2.18 million | 30.5% | 91.5%<br>(975/1066) | 6.0%<br>(64/1066) | 6.3 million<br>(3.9-10.0) | DRR347029-347052,<br>DRR347059-347074 |
| *C. monstifica* | 24 | 240 million | 2.29 million | 31.0% | 91.7%<br>(977/1066) | 5.9%<br>(63/1066) | 7.6 million<br>(5.2-13.3) | DRR346983-347006 |

**Table S5.** Summary of annotation for soldier or reproductive-biased transcripts among at least three *Colophina* species.

| Bias | Category | Over-represented<br>P value | GO term | Description | Species |
| --- | --- | --- | --- | --- | --- |
| Soldier | GO:0044449 | 2.75E-07 | contractile fiber part | CC | arma-clematis-monstrifica |
| Soldier | GO:0030018 | 5.41E-07 | Z disc | CC | arma-clematis-monstrifica |
| Soldier | GO:0005859 | 1.22E-05 | muscle myosin complex | CC | arma-clematis-monstrifica |
| Soldier | GO:0030017 | 2.37E-05 | sarcomere | CC | arma-clematis-monstrifica |
| Soldier | GO:0008307 | 5.85E-05 | structural constituent of muscle | MF | arma-clematis-monstrifica |
| Soldier | GO:0016460 | 8.32E-05 | myosin II complex | CC | arma-clematis-monstrifica |
| Soldier | GO:0005863 | 1.29E-04 | striated muscle myosin thick<br>filament | CC | arma-clematis-monstrifica |
| Soldier | GO:0032982 | 1.29E-04 | myosin filament | CC | arma-clematis-monstrifica |
| Soldier | GO:0062023 | 1.45E-04 | collagen-containing<br>extracellular matrix | CC | arma-clematis-monstrifica |
| Soldier | GO:0007517 | 3.16E-04 | muscle organ development | BP | arma-clematis-monstrifica |
| Soldier | GO:0045214 | 3.74E-04 | sarcomere organization | BP | arma-clematis-monstrifica |
| Soldier | GO:0044420 | 3.94E-04 | extracellular matrix component | CC | arma-clematis-monstrifica |
| Soldier | GO:0007520 | 5.66E-04 | myoblast fusion | BP | arma-clematis-monstrifica |
| Soldier | GO:0036379 | 6.88E-04 | myofilament | CC | arma-clematis-monstrifica |
| Soldier | GO:0000768 | 8.19E-04 | syncytium formation by plasma<br>membrane fusion | BP | arma-clematis-monstrifica |
| Soldier | GO:0006949 | 9.66E-04 | syncytium formation | BP | arma-clematis-monstrifica |
| Soldier | GO:0003779 | 3.28E-03 | actin binding | MF | arma-clematis-monstrifica |
| Soldier | GO:0005892 | 6.01E-03 | acetylcholine-gated channel<br>complex | CC | arma-clematis-monstrifica |
| Soldier | GO:0031589 | 6.18E-03 | cell-substrate adhesion | BP | arma-clematis-monstrifica |
| Soldier | GO:0007271 | 6.24E-03 | synaptic transmission,<br>cholinergic | BP | arma-clematis-monstrifica |
| Soldier | GO:0061061 | 6.37E-03 | muscle structure development | BP | arma-clematis-monstrifica |
| Soldier | GO:0007498 | 7.40E-03 | mesoderm development | BP | arma-clematis-monstrifica |
| Soldier | GO:0016459 | 8.64E-03 | myosin complex | CC | arma-clematis-monstrifica |
| Soldier | GO:0031032 | 9.78E-03 | actomyosin structure organization | BP | arma-clematis-monstrifica |
| Soldier | GO:0005509 | 9.98E-03 | calcium ion binding | MF | arma-clematis-monstrifica |
| Soldier | GO:0008092 | 1.33E-02 | cytoskeletal protein binding | MF | arma-clematis-monstrifica |
| Soldier | GO:0009888 | 1.59E-02 | tissue development | BP | arma-clematis-monstrifica |
| Soldier | GO:0030016 | 1.67E-02 | myofibril | CC | arma-clematis-monstrifica |
| Soldier | GO:0034446 | 1.88E-02 | substrate adhesion-dependent<br>cell spreading | BP | arma-clematis-monstrifica |
| Soldier | GO:0007155 | 2.02E-02 | cell adhesion | BP | arma-clematis-monstrifica |
| Soldier | GO:0043292 | 2.29E-02 | contractile fiber | CC | arma-clematis-monstrifica |
| Soldier | GO:0042805 | 2.38E-02 | actinin binding | MF | arma-clematis-monstrifica |
| Soldier | GO:0045211 | 2.53E-02 | postsynaptic membrane | CC | arma-clematis-monstrifica |
| Soldier | GO:0050839 | 2.75E-02 | cell adhesion molecule binding | MF | arma-clematis-monstrifica |
| Soldier | GO:0022610 | 2.89E-02 | biological adhesion | BP | arma-clematis-monstrifica |
| Soldier | GO:0048646 | 3.20E-02 | anatomical structure formation<br>involved in morphogenesis | BP | arma-clematis-monstrifica |
| Soldier | GO:0003008 | 3.39E-02 | system process | BP | arma-clematis-monstrifica |
| Soldier | GO:0008466 | 3.55E-02 | glycogenin glucosyltransferase<br>activity | MF | arma-clematis-monstrifica |
| Soldier | GO:0102751 | 3.55E-02 | UDP-alpha-D-glucose:glucosyl-<br>glycogenin<br>alpha-D-glucosyltransferase<br>activity | MF | arma-clematis-monstrifica |
| Soldier | GO:0035767 | 3.55E-02 | endothelial cell chemotaxis | BP | arma-clematis-monstrifica |
| Soldier | GO:0070278 | 3.55E-02 | extracellular matrix constituent<br>secretion | BP | arma-clematis-monstrifica |
| Soldier | GO:0014873 | 3.64E-02 | response to muscle activity<br>involved in regulation of muscle<br>adaptation | BP | arma-clematis-monstrifica |
| Soldier | GO:0044456 | 3.05E-17 | synapse part | CC | clematis-clematicola-monstrifica |
| Soldier | GO:0007268 | 5.14E-10 | chemical synaptic transmission | BP | clematis-clematicola-monstrifica |
| Soldier | GO:0023052 | 1.11E-09 | signaling | BP | clematis-clematicola-monstrifica |
| Soldier | GO:0007267 | 5.91E-09 | cell-cell signaling | BP | clematis-clematicola- monstrifica |
| Soldier | GO:0034702 | 1.48E-07 | ion channel complex | CC | clematis-clematicola-monstrifica |
| Soldier | GO:0007218 | 2.13E-05 | neuropeptide signaling pathway | BP | clematis-clematicola-monstrifica |
| Soldier | GO:0005576 | 7.10E-05 | extracellular region | CC | clematis-clematicola-monstrifica |
| Soldier | GO:0022848 | 1.06E-04 | acetylcholine-gated cation-selective channel activity | MF | clematis-clematicola-monstrifica |
| Soldier | GO:0022824 | 4.54E-04 | transmitter-gated ion channel activity | MF | clematis-clematicola-monstrifica |
| Soldier | GO:0022835 | 4.54E-04 | transmitter-gated channel activity | MF | clematis-clematicola-monstrifica |
| Soldier | GO:0007215 | 5.65E-03 | glutamate receptor signaling pathway | BP | clematis-clematicola-monstrifica |
| Soldier | GO:0016533 | 4.05E-02 | protein kinase 5 complex | CC | clematis-clematicola-monstrifica |
| Soldier | GO:0021722 | 4.05E-02 | superior olivary nucleus maturation | BP | clematis-clematicola-monstrifica |
| Reproductive | GO:0016491 | 8.65E-09 | oxidoreductase activity | MF | arma-clematis-monstrifica |
| Reproductive | GO:0016614 | 2.36E-07 | oxidoreductase activity, acting on CH-OH group of donors | MF | arma-clematis-monstrifica |
| Reproductive | GO:0016616 | 2.85E-07 | oxidoreductase activity, acting on the CH-OH group of donors, NAD or NADP as acceptor | MF | arma-clematis-monstrifica |
| Reproductive | GO:0006633 | 6.38E-07 | fatty acid biosynthetic process | BP | arma-clematis-monstrifica |
| Reproductive | GO:0016417 | 1.19E-06 | S-acyltransferase activity | MF | arma-clematis-monstrifica |
| Reproductive | GO:0072330 | 2.64E-06 | monocarboxylic acid biosynthetic process | BP | arma-clematis-monstrifica |
| Reproductive | GO:0071353 | 3.89E-06 | cellular response to interleukin-4 | BP | arma-clematis-monstrifica |
| Reproductive | GO:0032787 | 8.46E-06 | monocarboxylic acid metabolic process | BP | arma-clematis-monstrifica |
| Reproductive | GO:0016836 | 2.76E-05 | hydro-lyase activity | MF | arma-clematis-monstrifica |
| Reproductive | GO:0016788 | 3.52E-05 | hydrolase activity, acting on ester bonds | MF | arma-clematis-monstrifica |
| Reproductive | GO:0006790 | 9.90E-05 | sulfur compound metabolic process | BP | arma-clematis-monstrifica |
| Reproductive | GO:0016829 | 1.10E-04 | lyase activity | MF | arma-clematis-monstrifica |
| Reproductive | GO:0016835 | 1.29E-04 | carbon-oxygen lyase activity | MF | arma-clematis-monstrifica |
| Reproductive | GO:0016053 | 1.86E-04 | organic acid biosynthetic process | BP | arma-clematis-monstrifica |
| Reproductive | GO:0046394 | 1.86E-04 | carboxylic acid biosynthetic process | BP | arma-clematis-monstrifica |
| Reproductive | GO:0044421 | 2.57E-04 | extracellular region part | CC | arma-clematis-monstrifica |
| Reproductive | GO:0016407 | 3.35E-04 | acetyltransferase activity | MF | arma-clematis-monstrifica |
| Reproductive | GO:0005576 | 1.02E-03 | extracellular region | CC | arma-clematis-monstrifica |
| Reproductive | GO:0019752 | 1.12E-03 | carboxylic acid metabolic process | BP | arma-clematis-monstrifica |
| Reproductive | GO:0015926 | 2.02E-03 | glucosidase activity | MF | arma-clematis-monstrifica |
| Reproductive | GO:0016209 | 2.22E-03 | antioxidant activity | MF | arma-clematis-monstrifica |
| Reproductive | GO:0008610 | 2.26E-03 | lipid biosynthetic process | BP | arma-clematis-monstrifica |
| Reproductive | GO:0043436 | 2.40E-03 | oxoacid metabolic process | BP | arma-clematis-monstrifica |
| Reproductive | GO:0006082 | 2.77E-03 | organic acid metabolic process | BP | arma-clematis-monstrifica |
| Reproductive | GO:0010817 | 3.57E-03 | regulation of hormone levels | BP | arma-clematis-monstrifica |
| Reproductive | GO:0020037 | 3.59E-03 | heme binding | MF | arma-clematis-monstrifica |
| Reproductive | GO:0004518 | 3.65E-03 | nuclease activity | MF | arma-clematis-monstrifica |
| Reproductive | GO:0046906 | 4.76E-03 | tetrapyrrole binding | MF | arma-clematis-monstrifica |
| Reproductive | GO:0044283 | 4.95E-03 | small molecule biosynthetic process | BP | arma-clematis-monstrifica |
| Reproductive | GO:0071345 | 5.16E-03 | cellular response to cytokine stimulus | BP | arma-clematis-monstrifica |
| Reproductive | GO:0050661 | 5.19E-03 | NADP binding | MF | arma-clematis-monstrifica |
| Reproductive | GO:0042445 | 5.27E-03 | hormone metabolic process | BP | arma-clematis-monstrifica |
| Reproductive | GO:0005615 | 9.39E-03 | extracellular space | CC | arma-clematis-monstrifica |
| Reproductive | GO:0004601 | 1.02E-02 | peroxidase activity | MF | arma-clematis-monstrifica |
| Reproductive | GO:0016684 | 1.25E-02 | oxidoreductase activity, acting on peroxide as acceptor | MF | arma-clematis-monstrifica |
| Reproductive | GO:0034754 | 1.27E-02 | cellular hormone metabolic process | BP | arma-clematis-monstrifica |
| Reproductive | GO:0004521 | 1.49E-02 | endoribonuclease activity | MF | arma-clematis-monstrifica |
| Reproductive | GO:0016787 | 1.53E-02 | hydrolase activity | MF | arma-clematis-monstrifica |
| Reproductive | GO:0042744 | 1.82E-02 | hydrogen peroxide catabolic<br>process | BP | arma-clematis-monstrifica |
| Reproductive | GO:0003824 | 1.95E-02 | catalytic activity | MF | arma-clematis-monstrifica |
| Reproductive | GO:0050662 | 2.02E-02 | coenzyme binding | MF | arma-clematis-monstrifica |
| Reproductive | GO:0016860 | 2.05E-02 | intramolecular oxidoreductase<br>activity | MF | arma-clematis-monstrifica |
| Reproductive | GO:0019835 | 2.07E-02 | cytolysis | BP | arma-clematis-monstrifica |
| Reproductive | GO:0042743 | 2.21E-02 | hydrogen peroxide metabolic<br>process | BP | arma-clematis-monstrifica |
| Reproductive | GO:0044281 | 3.73E-02 | small molecule metabolic<br>process | BP | arma-clematis-monstrifica |
| Reproductive | GO:1901605 | 3.75E-02 | alpha-amino acid metabolic<br>process | BP | arma-clematis-monstrifica |
| Reproductive | GO:0090308 | 4.29E-02 | regulation of methylation-dependent chromatin silencing | BP | arma-clematis-monstrifica |
| Reproductive | GO:0090309 | 4.29E-02 | positive regulation of<br>methylation-dependent<br>chromatin silencing | BP | arma-clematis-monstrifica |
| Reproductive | GO:0048037 | 4.41E-02 | cofactor binding | MF | arma-clematis-monstrifica |
| Reproductive | GO:0035337 | 1.55E-05 | fatty-acyl-CoA metabolic<br>process | BP | arma-clematicola-monstrifica |
| Reproductive | GO:0016779 | 3.19E-04 | nucleotidyltransferase activity | MF | arma-clematicola-monstrifica |
| Reproductive | GO:0032450 | 4.70E-04 | maltose alpha-glucosidase<br>activity | MF | arma-clematicola-monstrifica |
| Reproductive | GO:0004558 | 6.64E-04 | alpha-1,4-glucosidase activity | MF | arma-clematicola-monstrifica |
| Reproductive | GO:0090599 | 1.21E-03 | alpha-glucosidase activity | MF | arma-clematicola-monstrifica |
| Reproductive | GO:0010025 | 1.46E-03 | wax biosynthetic process | BP | arma-clematicola-monstrifica |
| Reproductive | GO:0010166 | 1.46E-03 | wax metabolic process | BP | arma-clematicola-monstrifica |
| Reproductive | GO:0080019 | 1.46E-03 | fatty-acyl-CoA reductase<br>(alcohol-forming) activity | MF | arma-clematicola-monstrifica |
| Reproductive | GO:0102965 | 1.46E-03 | alcohol-forming fatty acyl-CoA<br>reductase activity | MF | arma-clematicola-monstrifica |
| Reproductive | GO:0006310 | 1.97E-03 | DNA recombination | BP | arma-clematicola-monstrifica |
| Reproductive | GO:0004553 | 2.57E-03 | hydrolase activity, hydrolyzing<br>O-glycosyl compounds | MF | arma-clematicola-monstrifica |
| Reproductive | GO:0005589 | 2.43E-02 | collagen type VI trimer | CC | arma-clematicola-monstrifica |
| Reproductive | GO:0016628 | 9.57E-08 | oxidoreductase activity, acting<br>on the CH-CH group of donors,<br>NAD or NADP as acceptor | MF | arma-clematis-clematicola |
| Reproductive | GO:0016627 | 6.44E-07 | oxidoreductase activity, acting<br>on the CH-CH group of donors | MF | arma-clematis-clematicola |
| Reproductive | GO:0042555 | 7.75E-06 | MCM complex | CC | clematis-clematicola-monstrifica |
| Reproductive | GO:0000776 | 9.36E-04 | kinetochore | CC | clematis-clematicola-monstrifica |
| Reproductive | GO:0006270 | 1.39E-03 | DNA replication initiation | BP | clematis-clematicola-monstrifica |
| Reproductive | GO:0006267 | 1.48E-03 | pre-replicative complex<br>assembly involved in nuclear<br>cell cycle DNA replication | BP | clematis-clematicola-monstrifica |
| Reproductive | GO:0036388 | 1.48E-03 | pre-replicative complex<br>assembly | BP | clematis-clematicola-monstrifica |
| Reproductive | GO:1902299 | 1.48E-03 | pre-replicative complex<br>assembly involved in cell cycle<br>DNA replication | BP | clematis-clematicola-monstrifica |
| Reproductive | GO:0000727 | 3.86E-03 | double-strand break repair via<br>break-induced replication | BP | clematis-clematicola-monstrifica |
| Reproductive | GO:0032133 | 4.60E-03 | chromosome passenger complex | CC | clematis-clematicola-monstrifica |
| Reproductive | GO:0006275 | 6.77E-03 | regulation of DNA replication | BP | clematis-clematicola-monstrifica |
| Reproductive | GO:0031618 | 8.75E-03 | nuclear pericentric<br>heterochromatin | CC | clematis-clematicola-monstrifica |
| Reproductive | GO:0005876 | 2.09E-02 | spindle microtubule | CC | clematis-clematicola-monstrifica |

BP, biological process; CC, cellular component; MF, molecular function

**Table S6.** Summary of annotation for transcripts consistently Soldier or Reproductive-biased among all four *Colophina* species.

| Bias | Category | Over-represented<br>P value | GO term | Description |
| --- | --- | --- | --- | --- |
| Reproductive | GO:0004313 | 2.52E-10 | [acyl-carrier-protein] S-acetyltransferase activity | MF |
| Reproductive | GO:0004320 | 2.52E-10 | oleoyl-[acyl-carrier-protein] hydrolase activity | MF |
| Reproductive | GO:0008659 | 2.52E-10 | (3R)-hydroxymyristoyl-[acyl-carrier-protein]<br>dehydratase activity | MF |
| Reproductive | GO:0016295 | 2.52E-10 | myristoyl-[acyl-carrier-protein] hydrolase activity | MF |
| Reproductive | GO:0016296 | 2.52E-10 | palmitoyl-[acyl-carrier-protein] hydrolase activity | MF |
| Reproductive | GO:0016297 | 2.52E-10 | acyl-[acyl-carrier-protein] hydrolase activity | MF |
| Reproductive | GO:0019171 | 2.52E-10 | 3-hydroxyacyl-[acyl-carrier-protein] dehydratase<br>activity | MF |
| Reproductive | GO:0047117 | 2.52E-10 | enoyl-[acyl-carrier-protein] reductase (NADPH,<br>A-specific) activity | MF |
| Reproductive | GO:0047451 | 2.52E-10 | 3-hydroxyoctanoyl-[acyl-carrier-protein]<br>dehydratase activity | MF |
| Reproductive | GO:0004314 | 5.41E-10 | [acyl-carrier-protein] S-malonyltransferase<br>activity | MF |
| Reproductive | GO:0016419 | 5.41E-10 | S-malonyltransferase activity | MF |
| Reproductive | GO:0016420 | 5.41E-10 | malonyltransferase activity | MF |
| Reproductive | GO:0004316 | 5.41E-10 | 3-oxoacyl-[acyl-carrier-protein] reductase<br>(NADPH) activity | MF |
| Reproductive | GO:0102131 | 5.41E-10 | 3-oxo-glutaryl-[acp] methyl ester reductase<br>activity | MF |
| Reproductive | GO:0102132 | 5.41E-10 | 3-oxo-pimeloyl-[acp] methyl ester reductase<br>activity | MF |
| Reproductive | GO:0004315 | 7.41E-10 | 3-oxoacyl-[acyl-carrier-protein] synthase activity | MF |
| Reproductive | GO:0031177 | 1.09E-09 | phosphopantetheine binding | MF |
| Reproductive | GO:0016418 | 2.67E-09 | S-acetyltransferase activity | MF |
| Reproductive | GO:0004312 | 1.95E-08 | fatty acid synthase activity | MF |
| Reproductive | GO:0008611 | 6.76E-06 | ether lipid biosynthetic process | BP |
| Reproductive | GO:0046504 | 6.76E-06 | glycerol ether biosynthetic process | BP |
| Reproductive | GO:1901503 | 6.76E-06 | ether biosynthetic process | BP |
| Reproductive | GO:0046485 | 9.25E-06 | ether lipid metabolic process | BP |
| Reproductive | GO:0004167 | 1.02E-05 | dopachrome isomerase activity | MF |
| Reproductive | GO:0050178 | 1.02E-05 | phenylpyruvate tautomerase activity | MF |
| Reproductive | GO:0070670 | 2.02E-05 | response to interleukin-4 | BP |
| Reproductive | GO:0016862 | 2.52E-05 | intramolecular oxidoreductase activity,<br>interconverting keto- and enol-groups | MF |
| Reproductive | GO:0006662 | 2.78E-05 | glycerol ether metabolic process | BP |
| Reproductive | GO:0018904 | 2.78E-05 | ether metabolic process | BP |
| Reproductive | GO:0097384 | 4.35E-05 | cellular lipid biosynthetic process | BP |
| Reproductive | GO:0016863 | 1.62E-04 | intramolecular oxidoreductase activity,<br>transposing C=C bonds | MF |
| Reproductive | GO:0004519 | 2.19E-03 | endonuclease activity | MF |
| Reproductive | GO:0005125 | 2.71E-03 | cytokine activity | MF |
| Reproductive | GO:0006259 | 4.47E-03 | DNA metabolic process | BP |
| Reproductive | GO:0016798 | 5.03E-03 | hydrolase activity, acting on glycosyl bonds | MF |
| Reproductive | GO:0007306 | 2.05E-02 | eggshell chorion assembly | BP |
| Soldier | GO:0022848 | 9.32E-03 | acetylcholine-gated cation-selective channel<br>activity | MF |
| Soldier | GO:0022824 | 1.41E-02 | transmitter-gated ion channel activity | MF |
| Soldier | GO:0022835 | 1.41E-02 | transmitter-gated channel activity | MF |

BP, biological process; CC, cellular component; MF, molecular function

**Table S7.** Correlation evolution among morphs. Shown are Pearson correlation coefficients calculated from phylogenetic independent contrasts in all transcripts (N = 18051) between a pair of morphs.

|  | gall1st | gall2nd | gallwingbud | gallwingless | opensol | opennor | openwingbud |
| --- | --- | --- | --- | --- | --- | --- | --- |
| gall1st |  |  |  |  |  |  |  |
| gall2nd | 0.77**** |  |  |  |  |  |  |
| gallwingbud | 0.64**** | 0.72**** |  |  |  |  |  |
| gallwingless | 0.81**** | 0.78**** | 0.72**** |  |  |  |  |
| opensol | 0.48**** | 0.49**** | 0.35**** | 0.44**** |  |  |  |
| opennor | 0.28**** | 0.33**** | 0.35**** | 0.31**** | 0.37**** |  |  |
| openwingbud | 0.20**** | 0.32**** | 0.39**** | 0.33**** | 0.30**** | 0.53**** |  |
| openwingless | 0.36**** | 0.45**** | 0.40**** | 0.45**** | 0.53**** | 0.53**** | 0.51**** |

\*\*\*\* $P < 0.0001$ .

**Table S8.** Correlation evolution among morphs. Shown are Pearson correlation coefficients calculated from phylogenetic independent contrasts in Soldier DET (N = 216) between a pair of morphs.

|  | gall1st | gall2nd | gallwingbud | gallwingless | opensol | opennor | openwingbud |
| --- | --- | --- | --- | --- | --- | --- | --- |
| gall1st |  |  |  |  |  |  |  |
| gall2nd | 0.76**** |  |  |  |  |  |  |
| gallwingbud | 0.27**** | 0.22** |  |  |  |  |  |
| gallwingless | 0.74**** | 0.58**** | 0.40**** |  |  |  |  |
| opensol | 0.52**** | 0.55**** | 0.26*** | 0.36**** |  |  |  |
| opennor | 0.28**** | 0.22** | 0.35**** | 0.20** | 0.18** |  |  |
| openwingbud | 0.29**** | 0.27**** | 0.13 | 0.20** | 0.15* | 0.75**** |  |
| openwingless | 0.17* | 0.13 | 0.42**** | 0.32**** | 0.33**** | 0.52**** | 0.53**** |

\*\*\*\* $P < 0.0001$ ; \*\*\* $P < 0.001$ ; \*\* $P < 0.01$ ; \* $P < 0.05$ .

**Table S9.** Predators of *Colophina clematis* found in their galls formed on *Zelkova serrata*.

| Collection locality<sup>a</sup> | Collection date | Prop. of galls containing predators |  |  |
| --- | --- | --- | --- | --- |
|  |  | Aphid flies<br>(Chamaemyiidae) | Hoverflies<br>(Syrphidae) | Lacewings<br>(Chrysapoidae) |
| Matsumoto, Nagano | July 17, 2019 | 49 % (22/45) | 2.2 % (1/45) | 2.2 % (1/45) |
| Okutama, Tokyo | June 19, 2018 | 50 % (13/16) | NA | NA |
| Shomaru, Saitama | July 4, 2018 | 60 % (3/5) | NA | NA |
| Total |  | 50<sup>b</sup> % (38/76) | 1.3<sup>c</sup> % (1/76) | 1.3<sup>c</sup> % (1/76) |

<sup>a</sup> All localities are in Japan.
<sup>bc</sup>Values within a column with different superscripts are significantly different (Fisher's exact tests, $P < 0.001$ ).

**Table S10.**
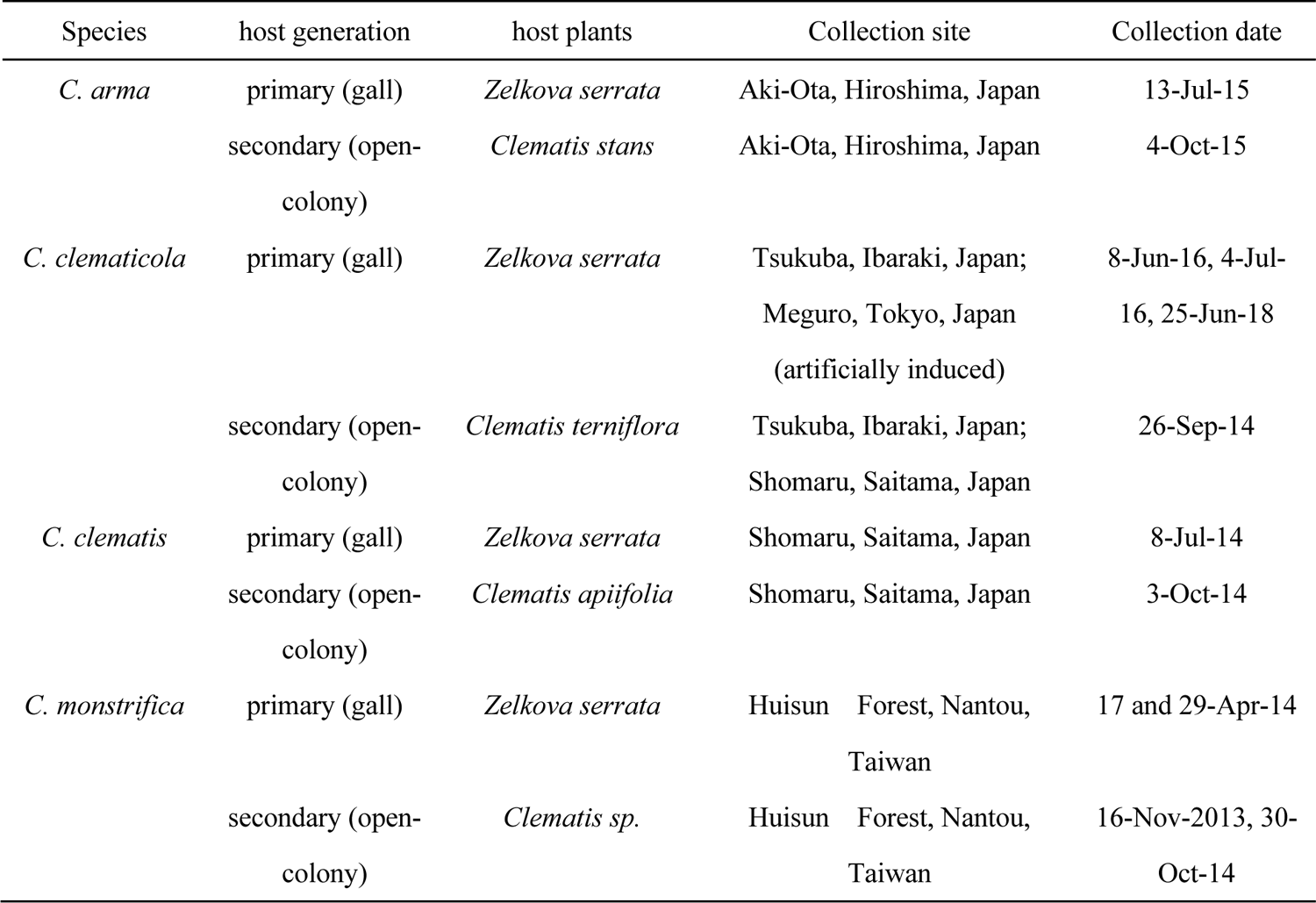
Collection sites of aphid species in this study.

## SI Figure legends

**Fig. S1.**
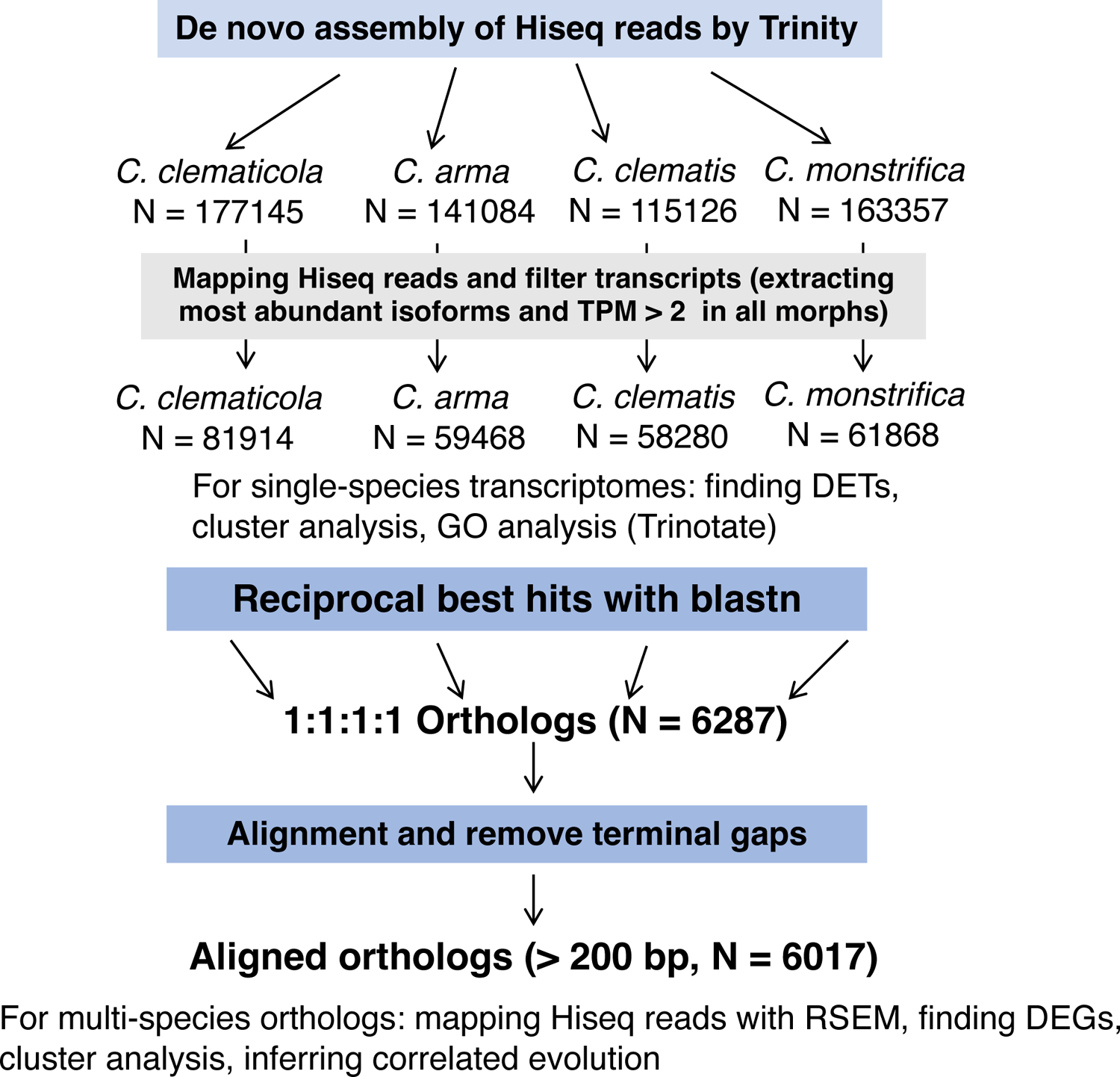
Workflow of transcriptome analysis.

**Fig. S2.**
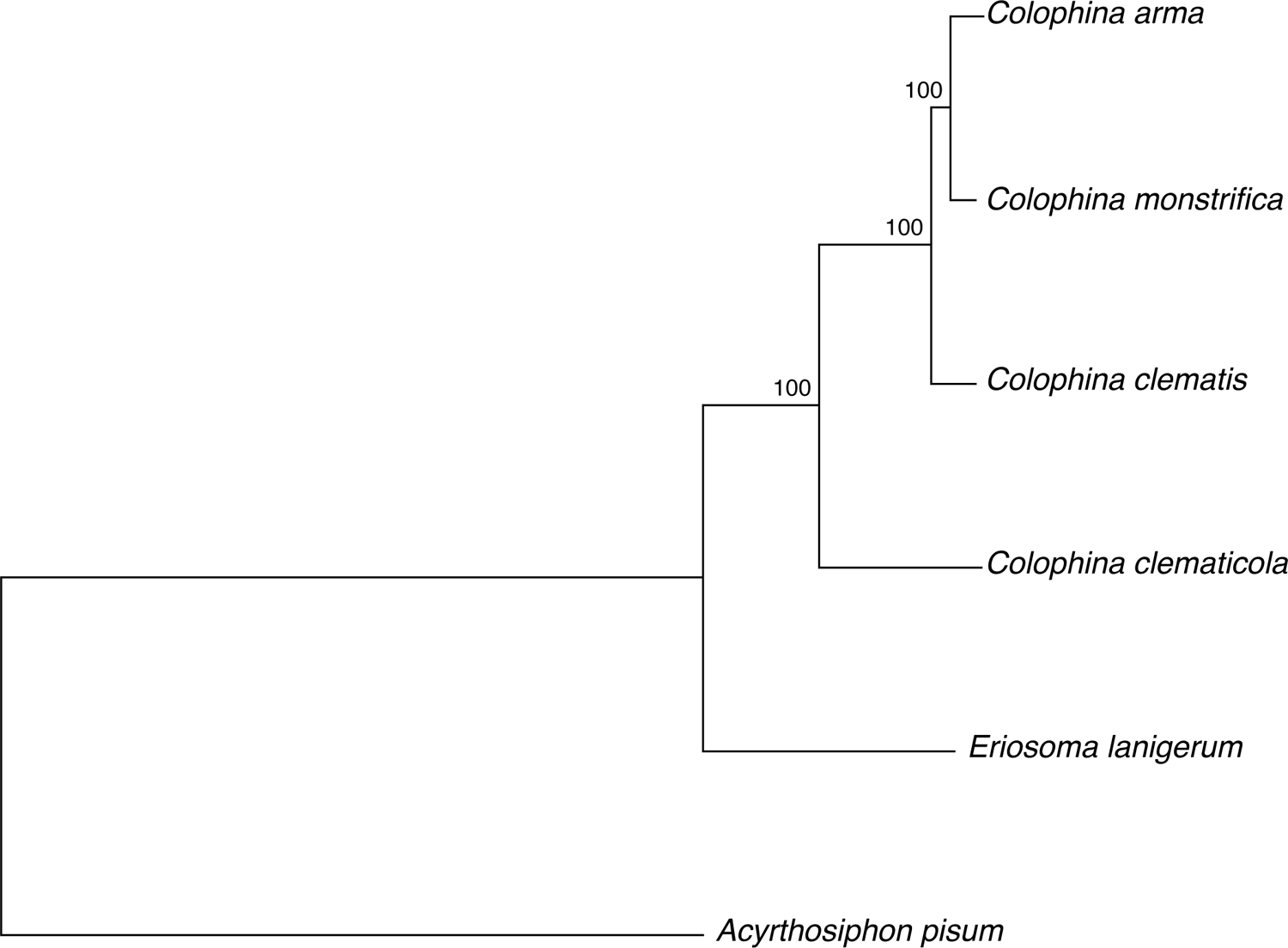
Molecular phylogenetic analysis of Eriosomatini aphids. A maximum-likelihood tree inferred from 3,755 one-to-one orthologous transcripts is shown. The pea aphid *Acyrthosiphon pisum* was used as an outgroup.

**Fig. S3.**
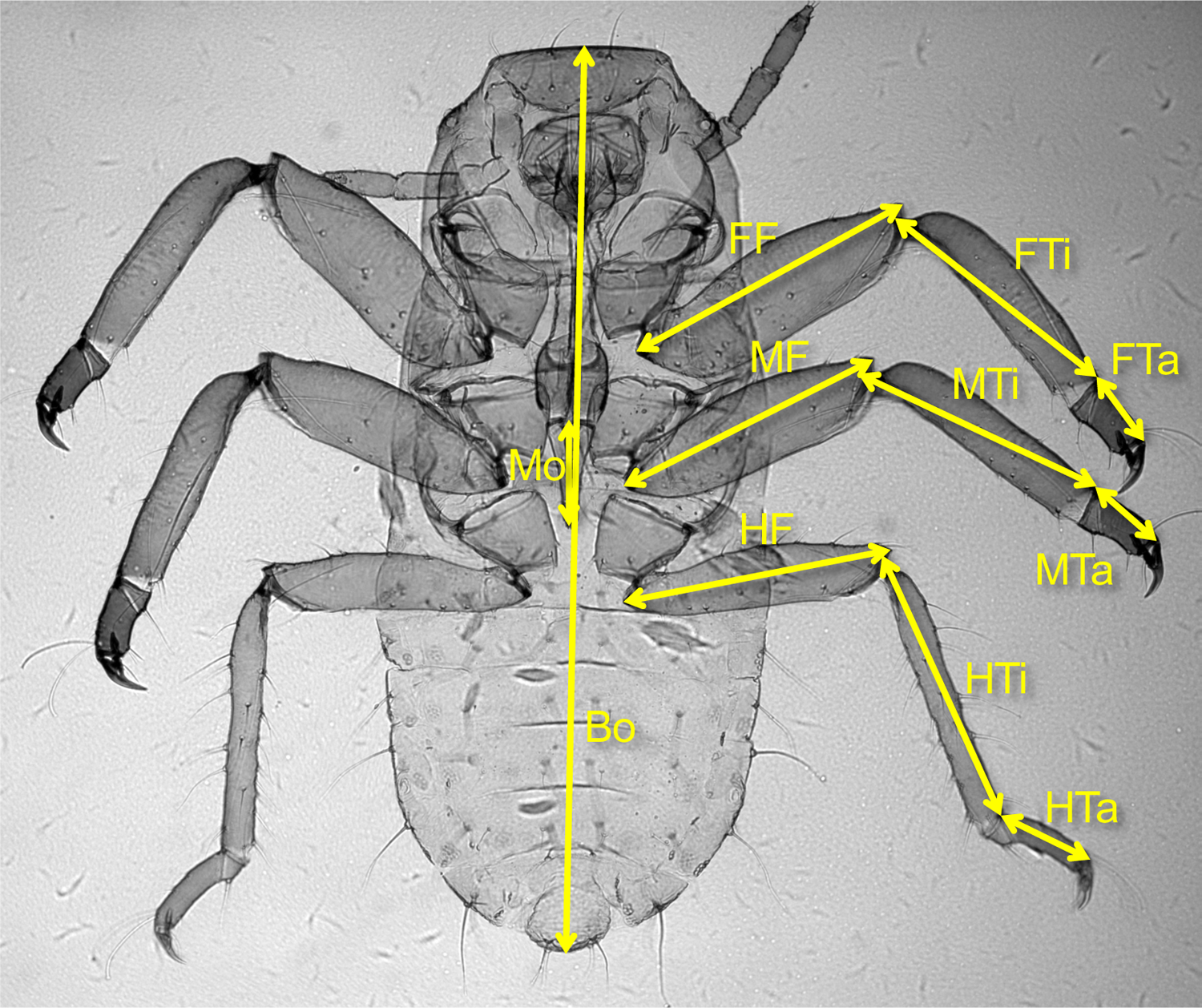
Morphological characters of aphids measured in this study. FFL, fore femur length; FTiL, fore tibia length; FTaL, fore tarsus length; MFL, mid femur length; MTiL, mid tibia length; MTaL, mid tarsus length; HFL, hind femur length; HTiL, hind tibia length; HTaL, hind tarsus length; ML, mouthpart length; BL, body length.

**Fig. S4.**
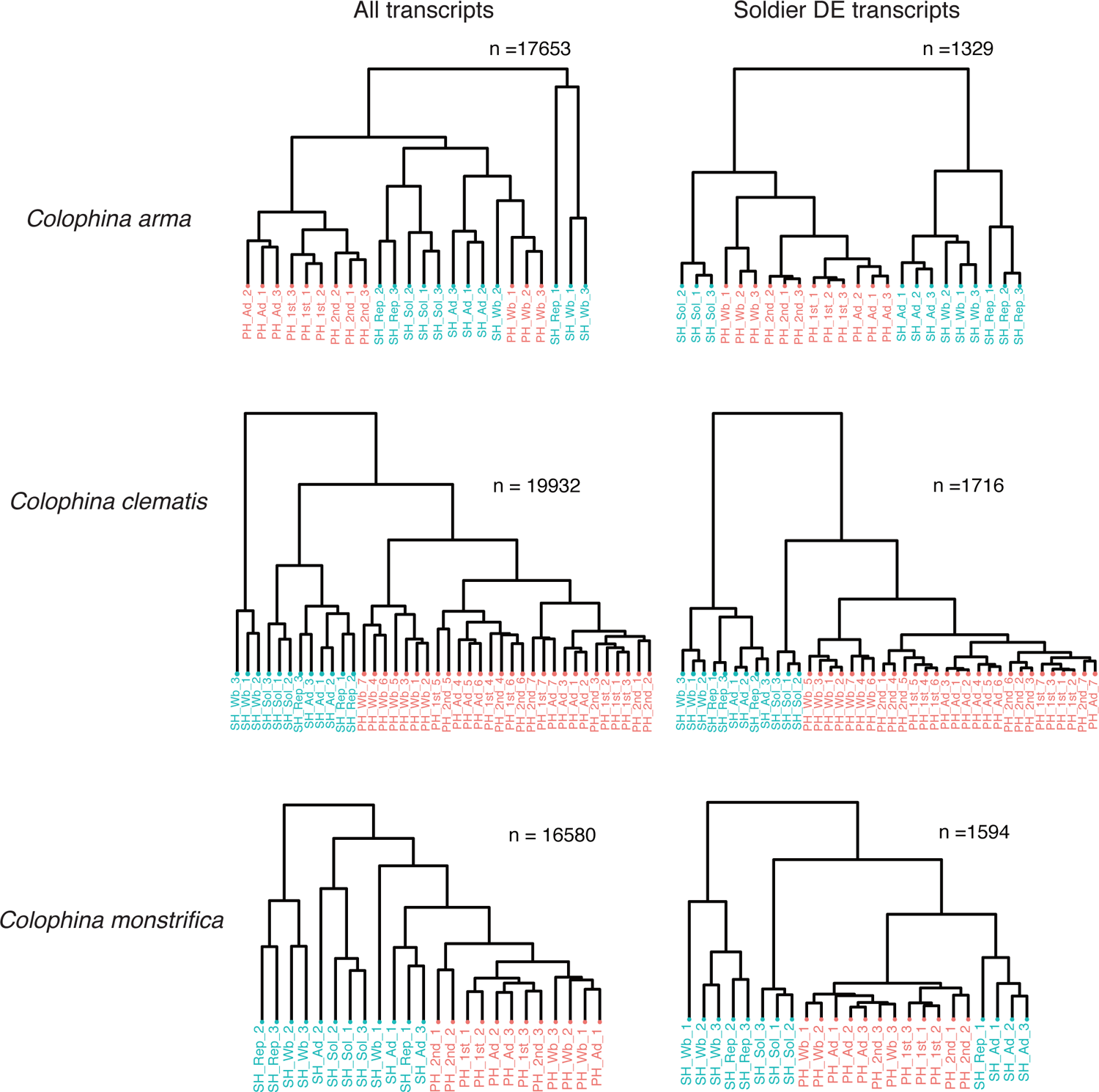
Single-species hierarchical clustering in transcriptome of *Colophina* aphids. Shown is the average relative expression including all transcripts (left) and Soldier differentially-expressed transcripts (DET, right). Hierarchical clustering of gene expression levels among four *Colophina* aphids using re-mapping raw reads to the aligned orthologous sequences of the one-to-one ortholog transcripts. Clustering is based on pairwise distance between samples computed as 1 - ρ, where ρ is Spearman’s correlation coefficient.

**Fig. S5.**
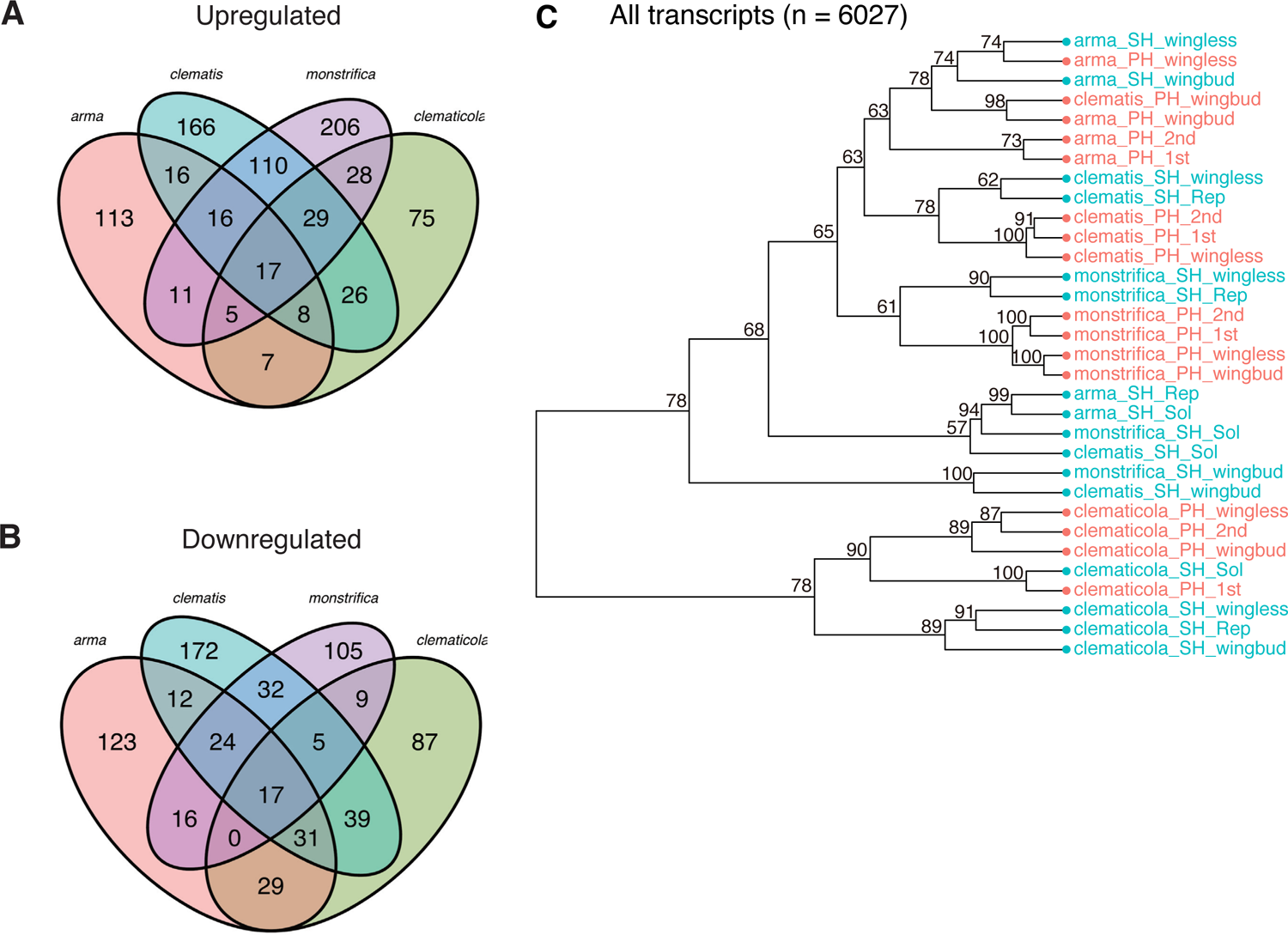
Venn diagram of the number of (A) differentially upregulated and (B) downregulated DET among 6,287 multispecies orthologous transcripts. The 17 upregulated and 17 downregulated overlapping transcripts across the four species with values that were 1119– and 686–fold higher than those predicted by chance (0.015 and 0.025; *P* < 0.001 for both cases; Fisher exact tests). (C) Hierarchical clustering of gene expression levels among four *Colophina* aphids using 6,017 orthologous transcripts across the four *Colophina* species. Colors denote host-plant generations: primary hosts (PH), red; secondary hosts (SH), blue. Abbreviations: 1st, first-instar nymph; 2nd, second-instar nymph; Wb, third- and fourth-instar nymph with wing buds; Ad, wingless adults.

**Fig. S6.**
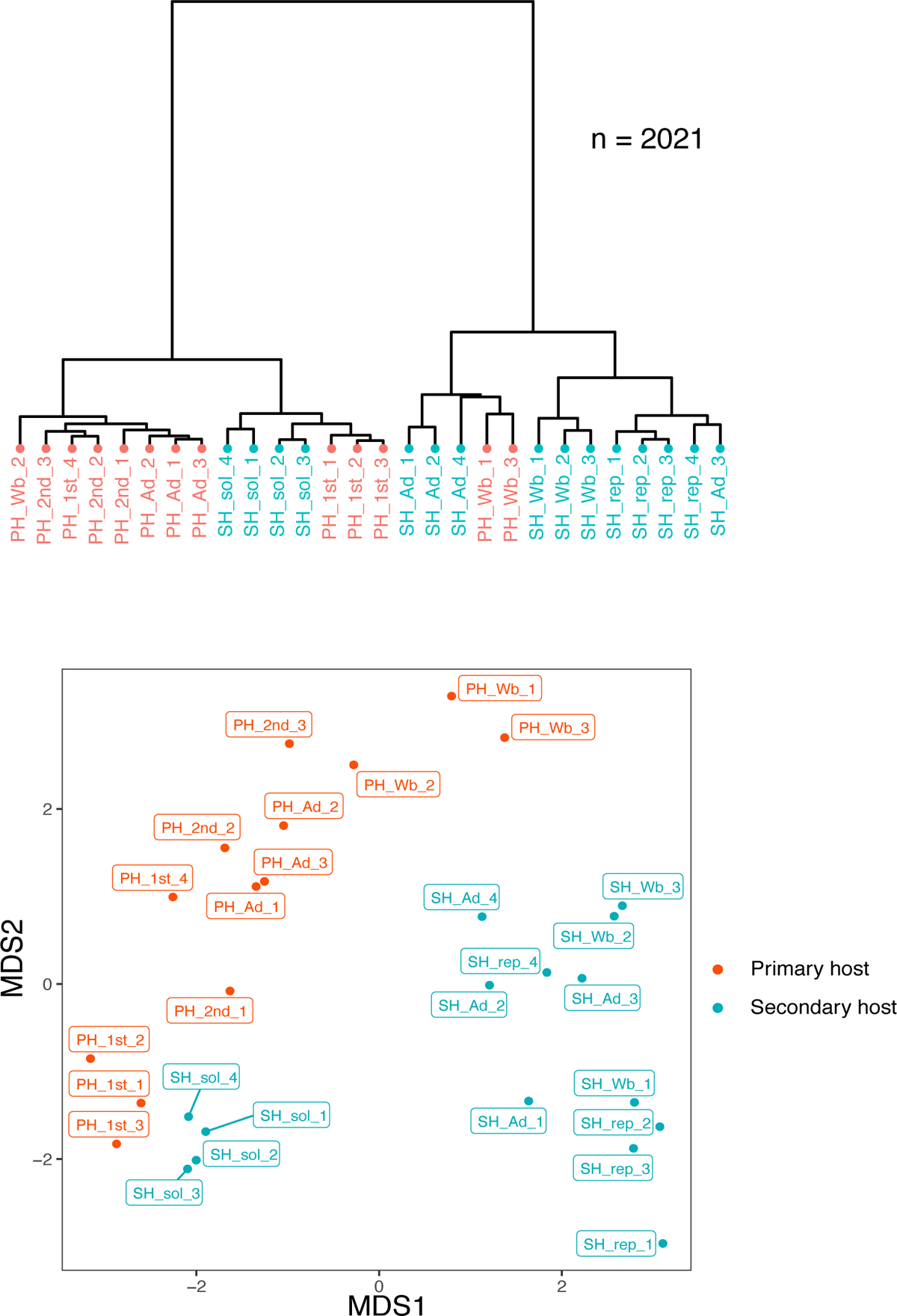
Similarities in transcriptomes between Gall defenders and Soldiers of *C. clematicola*. (A) Multidimensional Scaling (MDS) plot of the transcriptomes. (B) Hierarchical clustering of *Colophina clematicola* using Soldier DET (n = 2,021). Colors denote host-plant generations: primary hosts (PH), red; secondary hosts (SH).

**Fig. S7.**
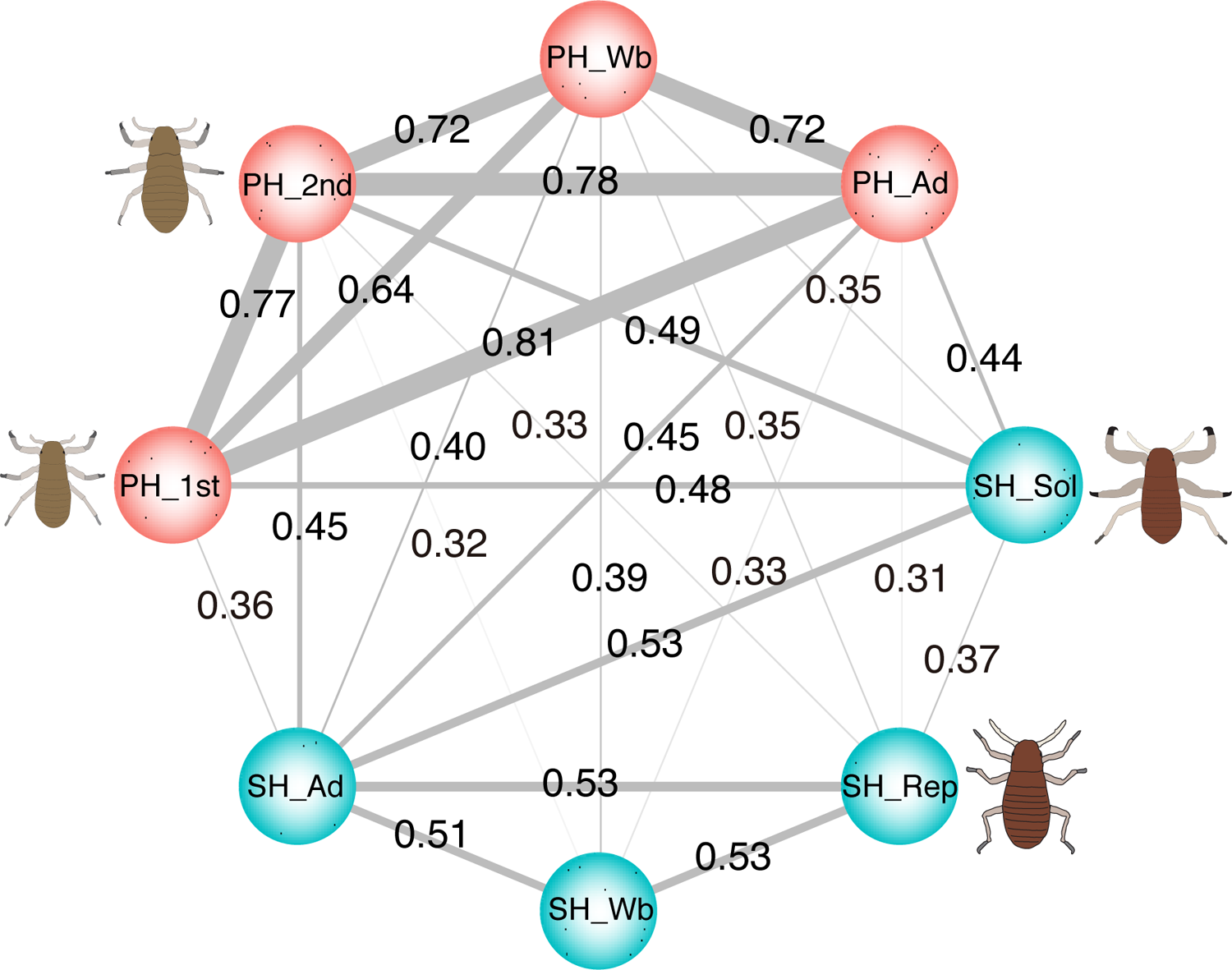
Correlation networks of the 6,017 orthologous transcripts. A number on the line represents Pearson’s Pairwise correlation coefficient (*r*) between a pair of morphs. Only *r* > 0.3 is shown as a line. Thicker line indicates a higher correlation. Colors denote host-plant generations: primary hosts (PH), red; secondary hosts (SH), blue. Abbreviations: 1st, first-instar nymph; 2nd, second-instar nymph; Wb, third- and fourth-instar nymph with wing buds; Ad, wingless adults.

**Fig. S8.**
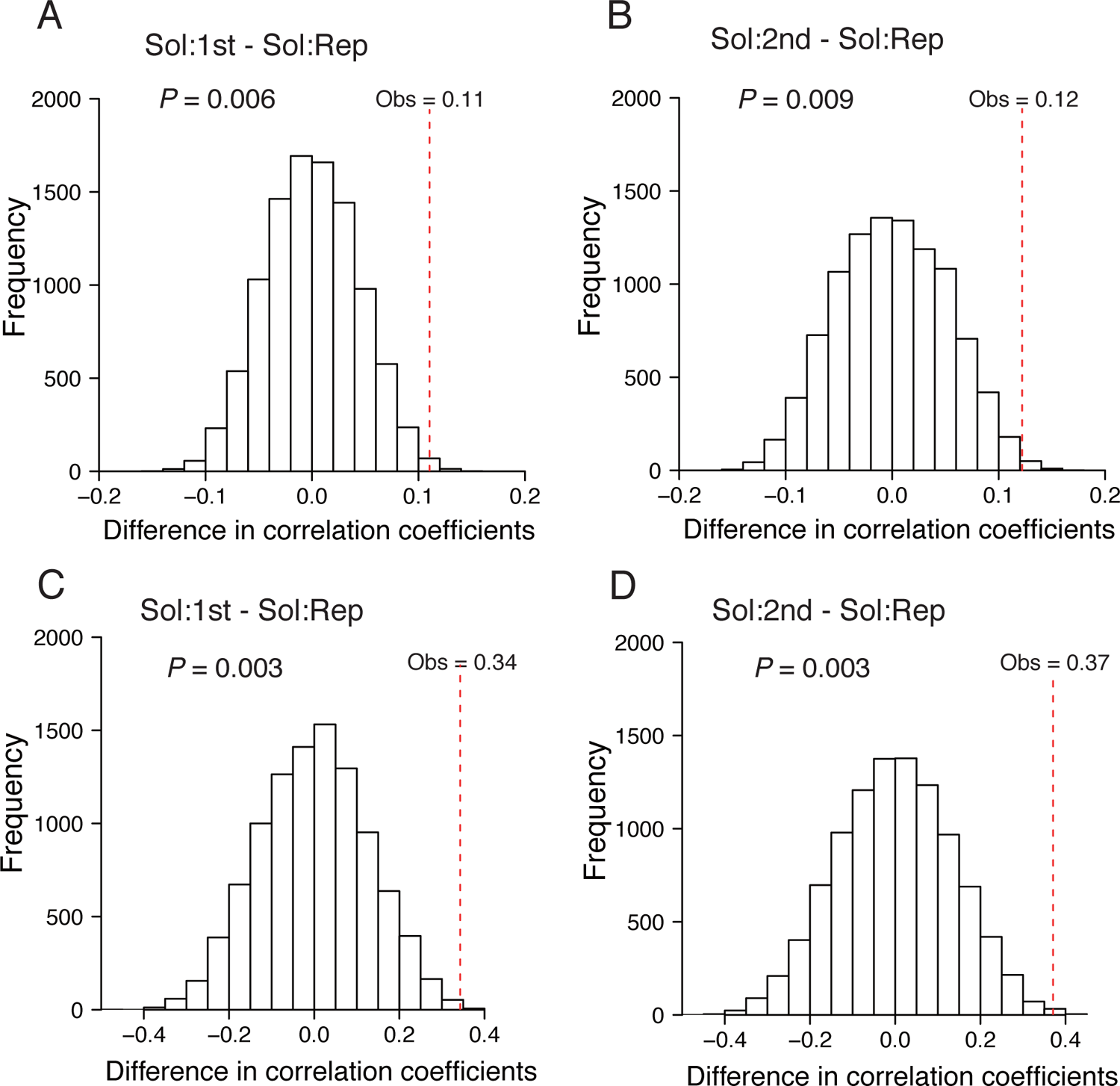
Permutation tests for significant difference in the degree of correlated evolution among morphs. Histograms show the distribution of difference in correlation coefficient for 10,000 permutated gene expression patterns, and the red dotted lines show the observed values of this difference between Soldier-Gall defender and Soldier-Reproductive correlation coefficients and for all orthologous transcripts and only Soldier DET. A, Soldier-Gall 1st-instar vs Soldier-Reproductive in all transcripts; B, Soldier-Gall 2nd-instar vs Soldier-Reproductive in all transcripts; C, Soldier-Gall 1st-instar vs Soldier-Reproductive in Soldier DET; D, Soldier-Gall 2nd-instar vs Soldier-Reproductive.

**Fig. S9.**
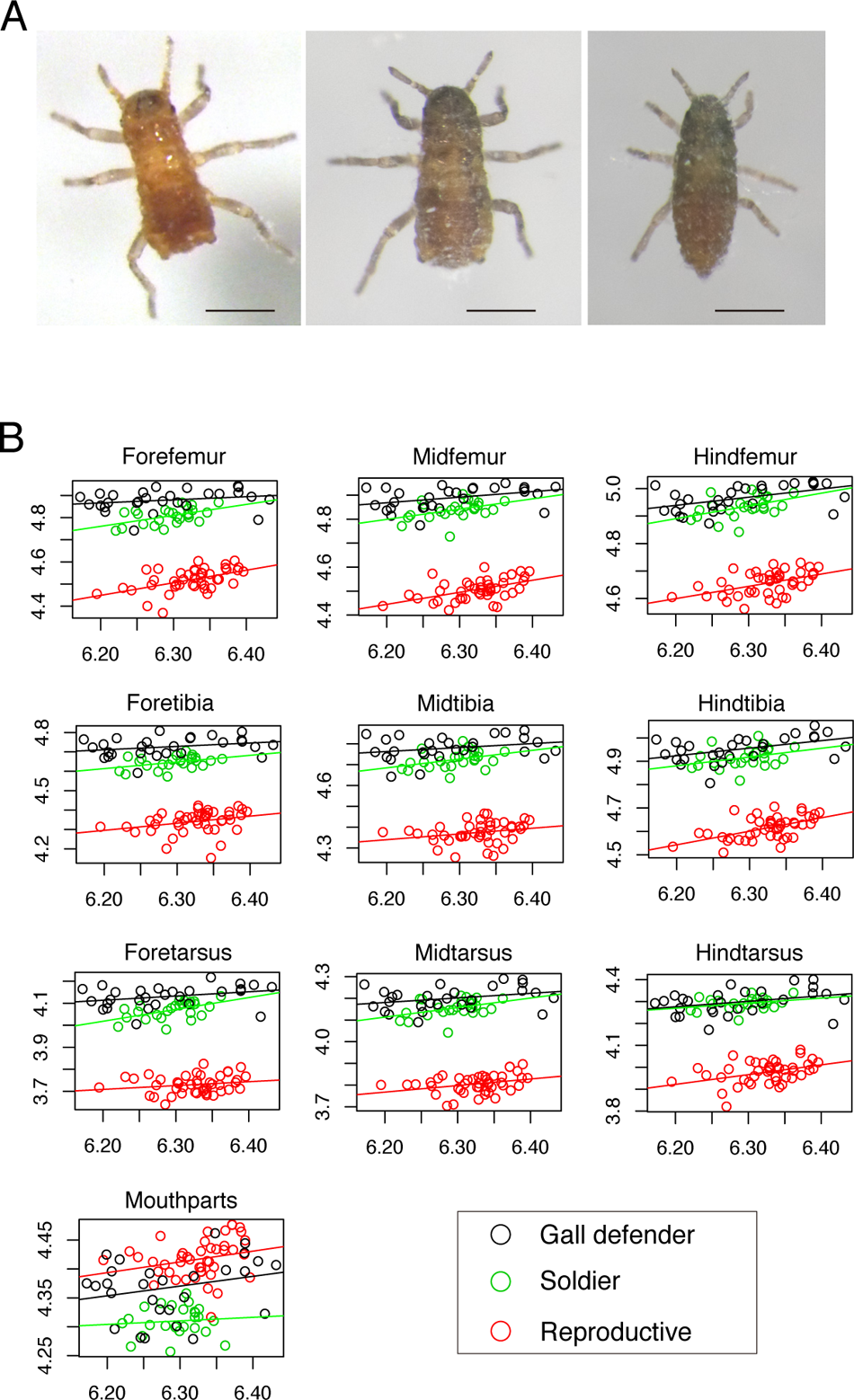
Similarities in morphology between Gall defenders and Soldiers of *C. clematicola*. (A) three morphs in the first-instar nymphs of *C. clematicola*: Gall defender on PH (Left); Soldier on SH (Middle); and Reproductive on SH (Right). Scale bars indicate 200 μm. (B) Distribution of morphological trait values among first instar nymphs in *C. clematicola*. Gall defenders: n = 28, shown in black; Soldiers: n = 25, shown in green; Reproductives: n = 44, shown in red. X-axis indicates log body size (μm) and y-axis log focal trait size (μm). ANCOVA results are shown in Table S2.

**Fig. S10.**
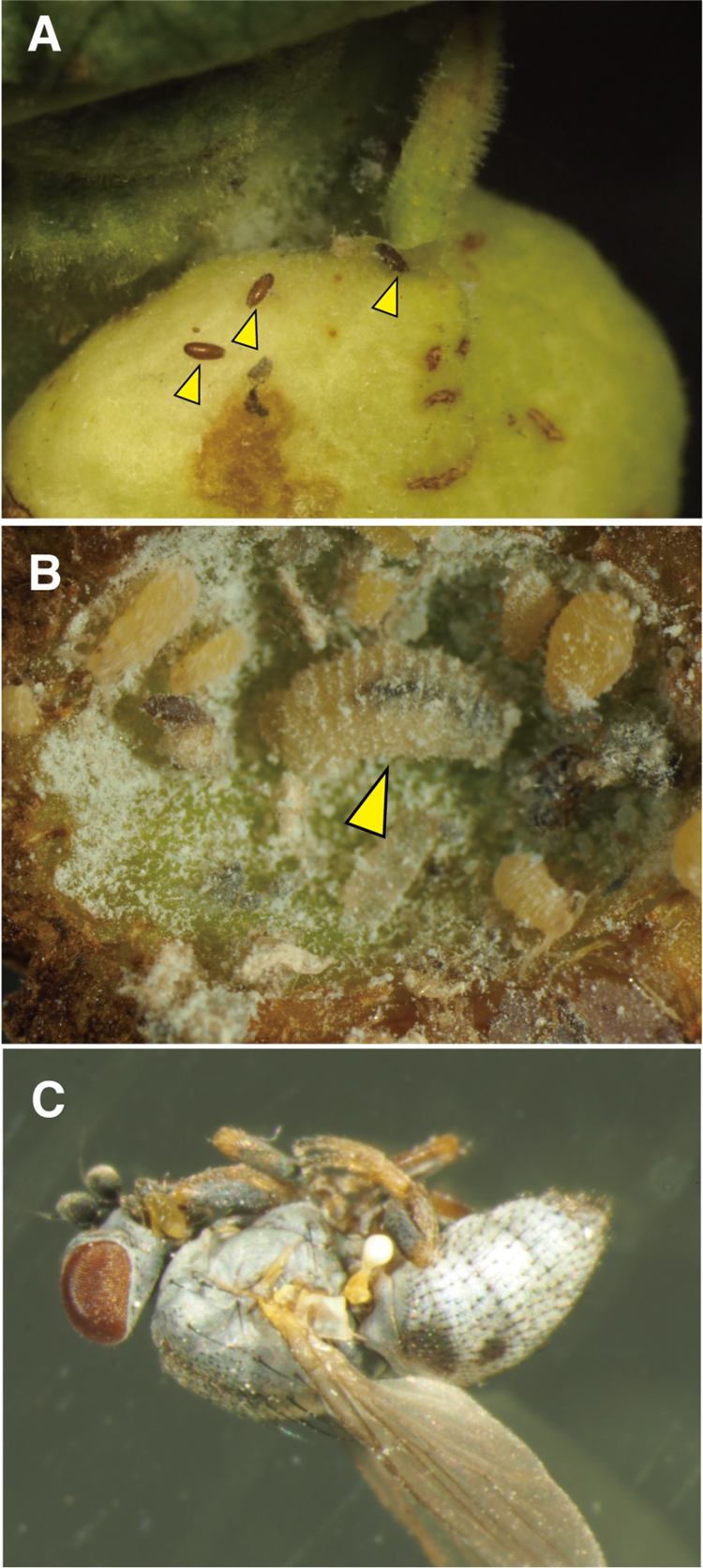
Predators of the primary host generation of *Colophina* aphids forming galls on *Zelkova serrata*. (A) Eggs of *Leucopis* sp. (arrowheads) found on the outer wall of a gall. (B) A larva of *Leucopis* sp. (arrowhead) predating in the gall of *C. clematis*. (C) An adult *Leucopis* fly emerged from a gall.

---

**Dataset S1**. Summary of caste differentially expressed transcripts in *Colophina arma*.

**Dataset S2**. Summary of caste differentially expressed transcripts in *Colophina clematis*.

**Dataset S3**. Summary of caste differentially expressed transcripts in *Colophina clematicola*.

**Dataset S4**. Summary of caste differentially expressed transcripts in *Colophina monstrifica*.

**Movie S1**. Attacking behavior of *C. arma* soldiers against a syrphid larva.

**Movie S2**. Attacking behavior of *C. clematicola* soldiers against a syrphid egg.

**Movie S3**. Attacking behavior of *C. arma* Gall defenders against a syrphid larva.

